# Pleomorphic effects of three small-molecule inhibitors on transcription elongation by *Mycobacterium tuberculosis* RNA polymerase

**DOI:** 10.1101/2025.02.07.637008

**Authors:** Omar Herrera-Asmat, Alexander B. Tong, Wenxia Lin, Tiantian Kong, Juan R Del Valle, Daniel G. Guerra, Yon W. Ebright, Richard H. Ebright, Carlos Bustamante

## Abstract

The *Mycobacterium tuberculosis* RNA polymerase (MtbRNAP) is the target of the first-line anti-tuberculosis inhibitor rifampin, however, the emergence of rifampin resistance necessitates the development of new antibiotics. Here, we communicate the first single-molecule characterization of MtbRNAP elongation and its inhibition by three diverse small-molecule inhibitors: N(α)-aroyl-N-aryl-phenylalaninamide (D-IX216), streptolydigin (Stl), and pseudouridimycin (PUM) using high-resolution optical tweezers. Compared to *Escherichia coli* RNA polymerase (EcoRNAP), MtbRNAP transcribes more slowly, has similar mechanical robustness, and only weakly recognizes *E. coli* pause sequences. The three small-molecule inhibitors of MtbRNAP exhibit strikingly different effects on transcription elongation. In the presence of D-IX216, which inhibits RNAP active-center bridge-helix motions required for nucleotide addition, the enzyme exhibits transitions between slowly and super-slowly elongating inhibited states. Stl, which inhibits the RNAP trigger-loop motions also required for nucleotide addition, inhibits RNAP primarily by inducing pausing and backtracking. PUM, a nucleoside analog of UTP, in addition to acting as a competitive inhibitor, induces the formation of slowly elongating RNAP inhibited states. Our results indicate that the three classes of small-molecule inhibitors affect the enzyme in distinct ways and show that the combination of Stl and D-IX216, which both target the RNAP bridge helix, has a strong synergistic effect on the enzyme.

## Introduction

Tuberculosis (TB) is the second most prevalent infectious disease causing morbidity and mortality worldwide (after COVID-19). There are 10 million new cases and 1.5 million deaths yearly, which incur $12 billion in medical and economic costs^1^. Rifampin is a small-molecule inhibitor of *Mycobacterium tuberculosis* RNA polymerase (MtbRNAP) and is a first-line treatment for TB^2,3^. The emergence and proliferation of rifampin resistance in *M. tuberculosis* have led to an effort to identify and develop new small-molecule inhibitors that effectively inhibit rifampin-resistant MtbRNAP^4–7^. A thorough understanding of the activity of MtbRNAP and its interactions with these new inhibitors is critical for informing and facilitating inhibitor development against TB.

In the last three decades, single-molecule optical tweezers studies of *E. coli* RNAP (EcoRNAP) and eukaryotic RNAP II have provided key insights on transcription elongation^8,9^, including the direct, real-time observation of single-base-pair stepping^10,11^. In addition, these studies have revealed details about sequence-dependent pausing, factor-dependent pausing^12,13^, and backtracking^13–16^. However, single-molecule studies of MtbRNAP and its inhibition by small-molecule inhibitors have been scarce. Current studies have primarily been structural and bulk assays^17–23^, with only one study by single-molecule methods addressing initiation^24^.

Here, we present a study of transcription elongation by MtbRNAP and its inhibition by small-molecule inhibitors using high-resolution single-molecule optical tweezers. First, we characterize *in singulo* the transcription elongation of MtbRNAP alone, including its pause-free velocity and mechanical robustness. Next, we assess three small-molecule inhibitors that target the transcription elongation of MtbRNAP. These inhibitors bind to sites distinct from rifampin and remain effective despite polymerase mutations that confer resistance to rifampin. These inhibitors are the N(α)-aroyl-N-aryl-phenylalaninamide (AAP) D-IX216^25,26^, streptolydigin (Stl)^27–32^, and pseudouridimycin (PUM)^33–35^.

AAPs selectively inhibit Mycobacterial RNAPs^26^ by binding to a site at the N-terminal end of the bridge helix. This site is conserved in Mycobacterial polymerases but not in other bacterial polymerases. The bound inhibitor interferes with the conformational dynamics of the bridge helix that are necessary for nucleotide addition^25,26,36,37^. We find that this inhibition results in the enzyme transitioning to different inhibition states that add nucleotides much more slowly. Streptolydigin (Stl) acts as a broad-spectrum inhibitor of bacterial RNAPs^27–29,31,32^. Stl binds to the region between the bridge helix and trigger loop, interfering with the closing of the trigger loop, which is required for nucleotide addition^29,32^. We find that Stl stabilizes pausing by the enzyme. PUM is another broad-spectrum inhibitor of bacterial RNAPs^33^ and is a nucleoside analog of UTP^33–35,38,39^, competitively inhibiting the binding of UTP. Additionally, PUM also appears to have an additional inhibitory effect on the enzyme^35^. Finally, when combined, D-IX216 and Stl exhibit synergistic inhibition of the enzyme.

## Results

### Biophysical and mechanochemical characterization of MtbRNAP

We studied the elongation of MtbRNAP using a high-resolution optical tweezers assay, where the RNAP and one end of the DNA template were tethered to polystyrene microspheres held in optical traps (Fig. 1a, left panel). We collected single-molecule trajectories of MtbRNAP transcription across a ∼3kb template with a sequence derived from *M. tuberculosis*.

**Figure 1:**
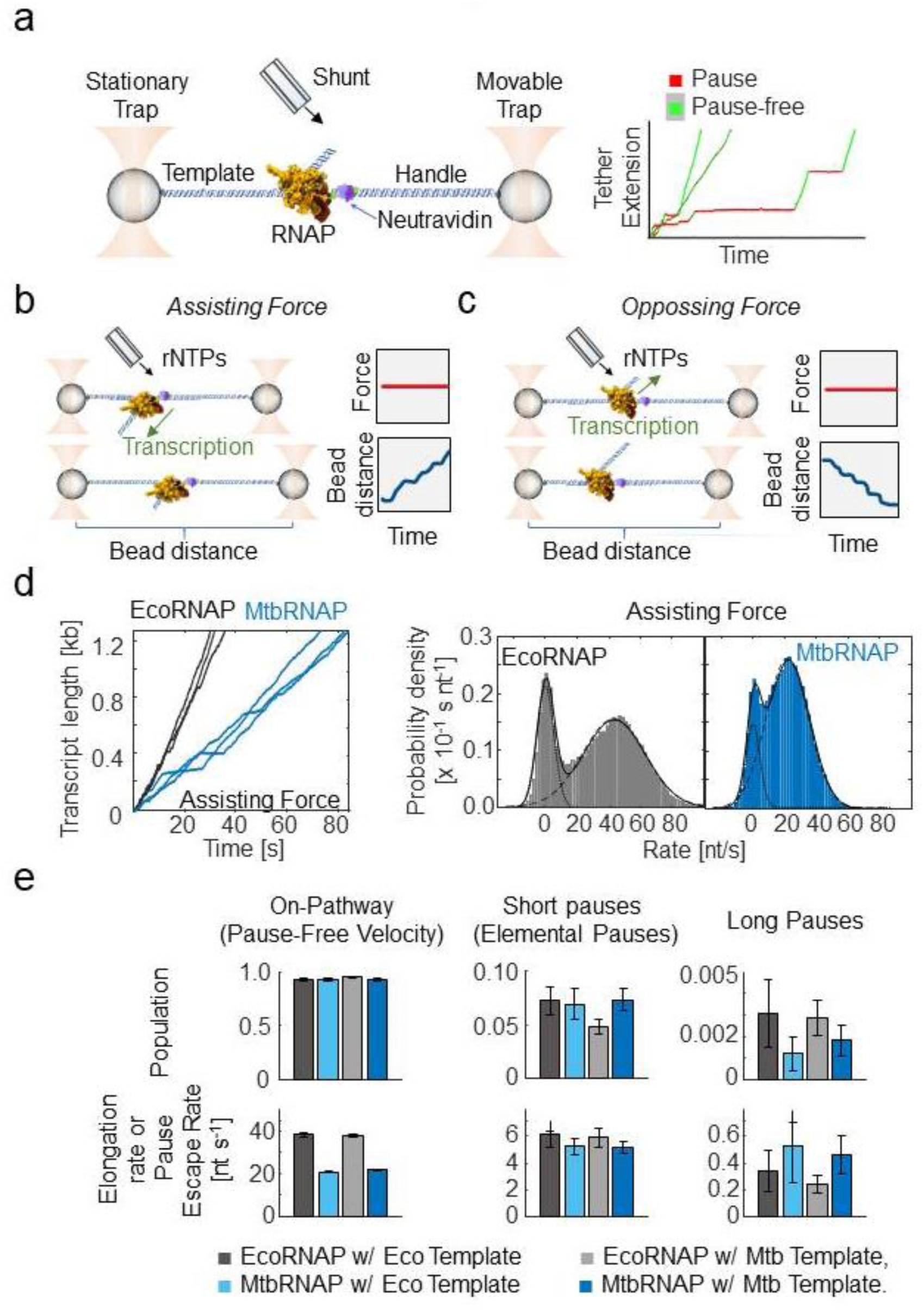
Single-molecule transcription elongation by *M. tuberculosis* RNA polymerase under high-resolution optical tweezers. a) (left) The experimental optical tweezer setup is shown, which tethers and isolates the RNAP from the laser beams through a DNA handle and template. (right) A few single-molecule transcription traces are shown, with the pauses (red) and the pause-free periods (green) highlighted. b) A diagram of assisting constant force mode is shown. The traps are moved as the RNAP transcribes to keep the force constant. c) A diagram of opposing constant force mode is shown. The traps are moved as the RNAP transcribes to keep the force constant. d) (left) Representative trajectories of transcription by EcoRNAP (black) and MtbRNAP (blue) under ∼18 pN constant assisting force, 22 °C, and saturating concentration of rNTPs (∼1mM) are shown. The DNA template for MtbRNAP derives from *M. tuberculosis* genes *rpoB* and *rpoC*. (right) Velocity distributions of these trajectories are shown, with a fit to a sum of two Gaussians, one with mean zero (pauses) and one with positive mean (pause-free velocity). e) The populations and rates obtained via DTD analysis are shown for EcoRNAP and MtbRNAP on Eco and Mtb templates. The traces were collected under saturating concentrations of nucleotides and 15-20 pN of assisting force for EcoRNAP or 18 pN for MtbRNAP.

Molecular trajectories were first obtained in *assisting force* mode (Fig.1b), where the upstream DNA is tethered, causing the force applied by the traps to aid the movement of the RNAP. We also obtained trajectories in *opposing force* mode (Fig.1c), where the downstream DNA is tethered, and the force hinders the enzyme’s movement. In both cases, as the RNAP moves, the length of the tether changes, and the position of the movable trap is adjusted to maintain a constant tension on the tether.

Transcription by MtbRNAP under an assisting force of 18 pN comprises periods of continuous activity at a consistent velocity interspersed with long pauses, a behavior also described for EcoRNAP^40–42^ (Fig. 1a, right panel). Plots of the velocity distribution of all molecular trajectories for both enzymes show a population at zero velocity corresponding to transcriptional pauses, and another corresponding to the pause-free velocity of the enzyme (Fig. 1d, right panel). The pause-free velocity of MtbRNAP is measured by this method is 23.1 ± 5.1 nt/s, approximately half that of EcoRNAP, which is 42.7 ± 8.6 nt/s (Fig. 1d, right panel).

To get a more precise characterization of the dynamics of RNAP elongation, we obtained the dwell time at each nucleotide by fitting the molecular traces to a monotonic one base-pair (bp) staircase function. We then modeled the resulting dwell time distribution (DTD) as a sum of exponentials (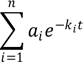)^43,44^. We interpret these different exponentials as distinct states of the enzyme: *a_i_* represents the state probability (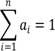), while *k_i_* indicates the kinetic rate of the *ith* state, such that *k_1_* > *k_2_*> … *k_n_*. The fastest state typically corresponds to the pause-free state, whereas the slower states reflect various dwell times associated with the paused states of the enzyme, including the relatively short elemental pauses and longer backtracking pauses.

For EcoRNAP, three exponential components effectively describe the DTD. The majority of the steps (92.4% ± 1.3%, 95% CI), belong to the faster state, with a rate of 38.3 ± 1.0 nt/s (*k_pause-free_*[*k_pf_*], 95% CI), which corresponds to the average rate of completing a pause-free nucleotide addition cycle^43^. The middle population, comprising 7.2% ± 1.3% (95% CI) of steps corresponded to a slower rate of 6.2 ± 1.0 nt/s (*k_elemental-pause_* [*k_ep_*], 95% CI), which we associate with the rate of escape by RNAP from elemental pauses^45^. Finally, the remaining 0.31% ± 0.16% of steps (95% CI) corresponds to an even slower rate of 0.34 ± 0.15 nt/s (95% CI), reflecting sporadic long pauses (*k_lp_*).

For MtbRNAP, we also identified three populations with a *k_pf_* of 20.7 ± 0.4 nt/s, a *k_ep_* 5.2 ± 0.6 s^-1^, and a *k_lp_* of 0.52 s^-1^ with similar population percentages to EcoRNAP (Fig. 1e). The pause-free velocity values obtained from the velocity distribution (Fig. 1d, right panel) and DTD (Fig. 1e) coincide well. We repeated this characterization for MtbRNAP under various magnitudes of assisting forces, ranging from 8 pN to 18 pN, and observed that *k_pf_* slightly increases with greater force, while *k_el_* was mostly unchanged. Conversely, when conducting the experiment under opposing forces from -4 to -18 pN, there are no significant effects on *k_pf_* or *k_el_*(Fig. 2a).

**Figure 2:**
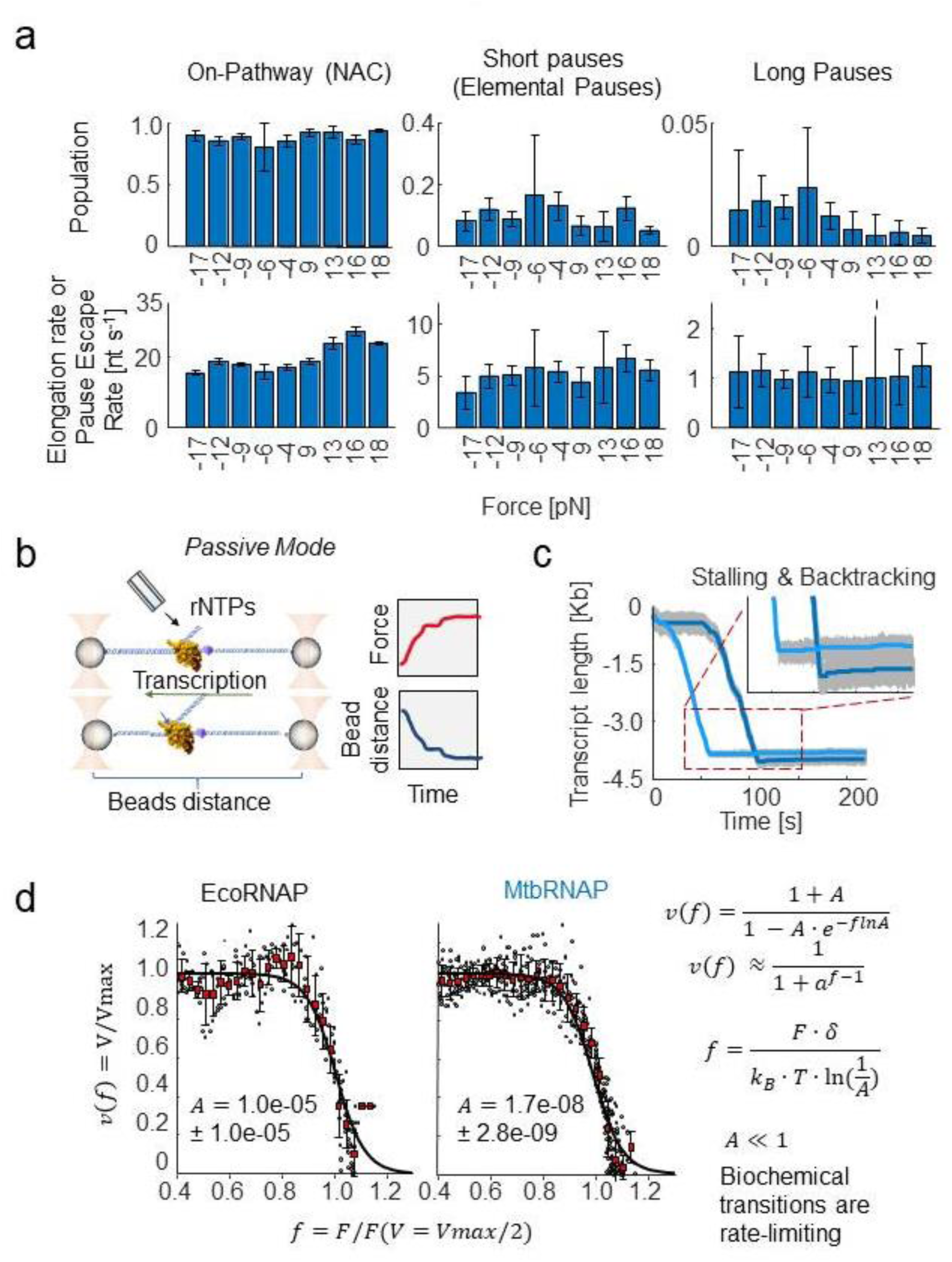
Mechanochemical characterization of the elongation by *M. tuberculosis* RNA polymerase under high-resolution optical tweezers. a) The effect of magnitude and direction of applied force on the kinetic parameters obtained from DTD analysis is shown. Positive forces are assisting forces, negative are opposing forces. The assisting force geometry moderately affects MtbRNAP’s effective elongation rate but not pause probability, while opposing forces increase the probability of long pauses. b) A diagram of passive opposing force mode is shown. The traps are held stationary as the RNAP transcribes, leading to an increase in force. c) Example passive mode traces are shown. The MtbRNAP transcribes at near constant velocity before suddenly stalling and backtracking. d) Force-velocity relationships for MtbRNAP and EcoRNAP obtained from passive mode experiments are shown and fit to a generalized Boltzmann relation (right). Both polymerases’ velocities remain insensitive to force until they reach similar stall forces, on average 19.3 pN ± 1.2 pN for MtbRNAP, and 17.1 ± 1.1 pN for EcoRNAP.

Next, we performed experiments in *passive mode*, where the downstream DNA is tethered in opposing force mode while the traps’ positions remain stationary, leading to an increase in force as transcription progresses (Fig. 2b). Passive mode is used to determine the stalling behavior and stall force of the polymerase. At low forces, MtbRNAP transcription velocity exhibits minimal response to the opposing force until a maximum force (*F_stall_*) is reached, beyond which the enzyme drastically slows, stalls, and backtracks (Fig. 2b)^45^. On average, MtbRNAP stalled at 19.3 ± 1.2 pN (s.e.m.)^46^. This force-induced stalling is often accompanied by backtracking (Fig. 2c). By comparison, EcoRNAP stalls at 17.1 ± 1.1 pN (s.e.m.) and is also accompanied by backtracking, consistent with previous studies^9,16,49^. The force-velocity curves can be fit to an unrestrained Boltzmann relation, where the process is split into biochemical (force-independent) and mechanical (force-dependent) terms, with their ratio governed by a parameter A (Fig. 2d)^11^.

For both Eco and Mtb RNAPs, we find that A << 1, indicating that biochemical transitions are rate-limiting in these instances, in line with prior findings for EcoRNAP^9^. This relation also includes the distance to the transition state δ, which dictates the force dependence of the mechanical term. We interpret this value as the degree of backtracking required by MtbRNAP to stall^47^. For MtbRNAP, we obtain a value of 14 ± 1 nt (95% CI) (Fig. 2d), comparable to the 13 ± 1 nt observed for EcoRNAP and findings from earlier studies on EcoRNAP^9^.

### MtbRNAP weakly recognizes pause sequences of EcoRNAP

We have observed that MtbRNAP tends to pause less frequently than EcoRNAP under an assisting force on both the Eco and Mtb templates (Fig. 1e). To gain a clearer understanding of the pausing dynamics, we utilized a *molecular ruler* template to compare the pausing behavior in a sequence-dependent manner. A molecular ruler consists of a tandem repeat of a sequence harboring one or more strong pauses (Fig. 3a)^48^. Because the trajectories of the different traces obtained on our high-resolution optical tweezers instrument all display the repeating pattern of pauses, it is possible to align them and determine the precise position of the RNAP on the template with nearly single base pair resolution^13,49^. Without a ruler, uncertainties in position caused by bead size variations or calibration errors cause the location of RNAP on the template to be not precisely known. We previously developed a ruler for EcoRNAP featuring five pauses designated as *a*, *b*, *c*, *d*, and *his*. However, we discovered that this ruler was ineffective for MtbRNAP, which does not pause strongly enough at those locations, as seen in the heights of the peaks of the residence time histogram (RTH) of the repeat (Fig. 3b, c). We confirmed this behavior using a gel assay, which shows weaker pausing by MtbRNAP at these locations (Fig. 3d). Notably, MtbRNAP demonstrates reduced recognition of the elemental pauses *a*, *b*, and *d*, and surprisingly did not strongly recognize the strong hairpin pause sequence *his* from *E. coli*. This last phenomenon could be attributed to differences in the interactions within the RNAP exit tunnel and the nascent RNA between the two enzymes^50^.

**Figure 3:**
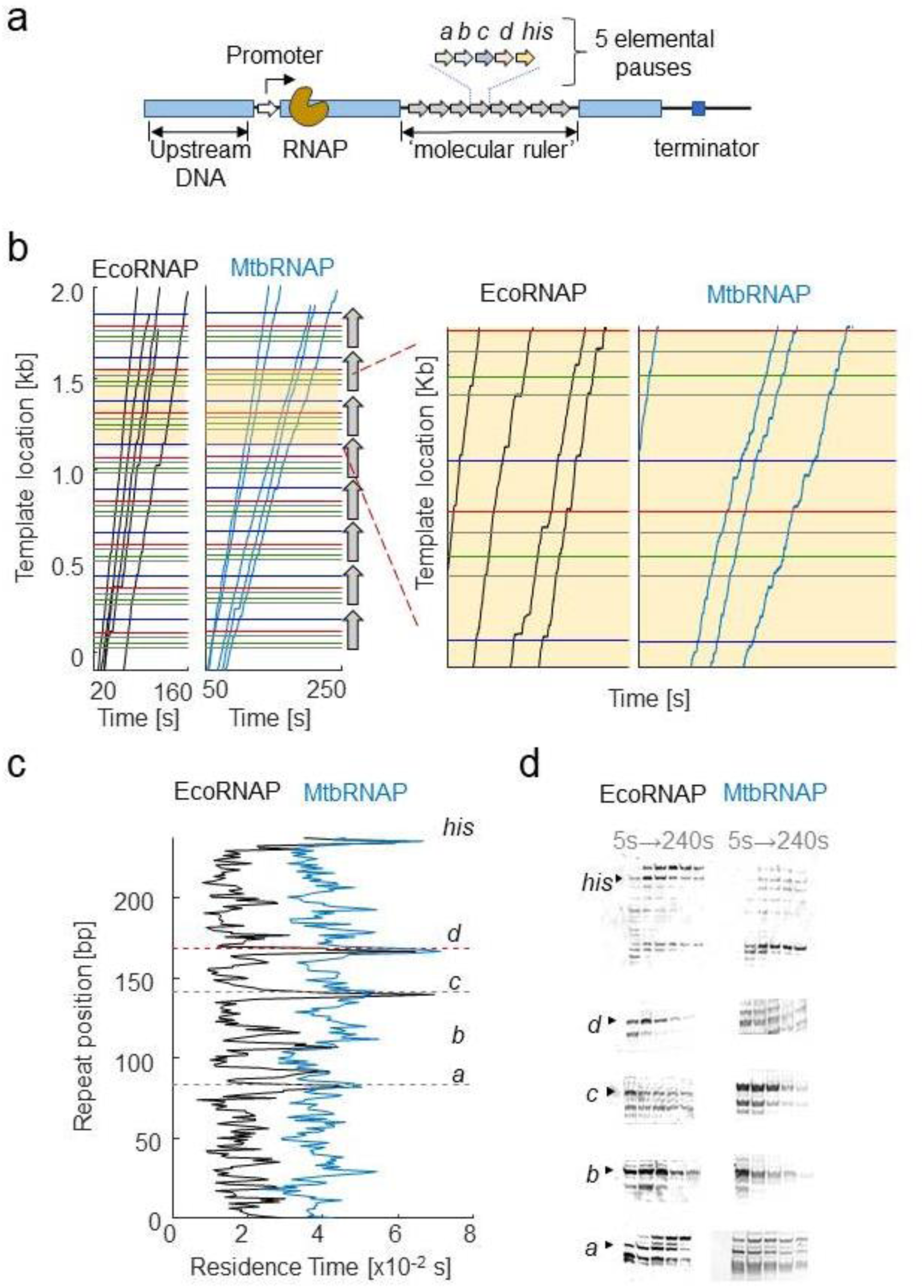
MtbRNAP pauses less efficiently than EcoRNAP on Eco pauses. a) Representation of a template containing a molecular ruler to study EcoRNAP pauses. The DNA template consists of 8 repeats of five *Eco* elementary pauses, *a, b*, *c*, *d*, and *his* arranged in tandem. b) Example traces of single Eco and Mtb RNA polymerases transcribing the *E. coli* molecular ruler are shown. The eight repeats are denoted as vertical arrows, and the location of the pauses within each repeat are shown as horizontal colored lines. We collected the transcription activities using a cocktail of rNTPs (1 mM rUTP, 1 mM rGTP, 0.5 mM rATP, and 0.25 mM rCTP) at 15-20 pN for EcoRNAP and 18 pN for MtbRNAP. The zoom on the right shows the alignment of the trace pauses with the expected pause locations. c) Residence time histograms (RTHs) of EcoRNAP and MtbRNAP display distinctive patterns of pause strength (peak height) and location. MtbRNAP more weakly recognizes the Eco pauses. d) Comparison of the transcription gel bands at the elemental pause sites between MtbRNAP and EcoRNAP. Figure 3—Source Data 1: Original gels for Figure 3d, indicating the conditions and relevant bands Figure 3—Source Data 2: Original files of the gels in Figure 3d.

Since MtbRNAP recognized pause *c* the strongest (Fig. 3d), we designed a new molecular ruler based on this sequence. The *Mtb* molecular ruler consists of eight repeats of the elemental pause *c,* inserted within a fragment of the *M. tuberculosis rpoB* gene. MtbRNAP pauses strongly at the expected location of 50 nt into the 68 nt repeat that makes up the ruler, with minor pauses at position 5 and 30 nt in the repeat region (Fig. S1). Using this tool, we proceeded to study the effects of the inhibitors in a sequence-dependent manner.

### D-IX216 inhibits elongation by MtbRNAP by switching it into interchangeable slow and super-slow inhibited states of transcription elongation

We investigated the effects of D-IX216 on the elongation activity of MtbRNAP. D-IX216 is an analog of D-AAP1^26^, having a 2-thiophenyl group in place of a phenyl group as ring A and a 5-fluoro-2-piperazin-1-yl-phenyl instead of 2-methyl-phenyl as ring C (Fig. S2a). This AAP analog is ten times more potent (IC_50_ of 100 nM determined via bulk transcription methods) against MtbRNAP compared to D-AAP1 (IC_50_ of 1µM)^25,26^, and is selective for MtbRNAP, not affecting EcoRNAP at these concentrations (Fig. S2b, c, d).

Upon adding 70-280 nM D-IX216 to a transcribing MtbRNAP (Fig. 4a), the elongation trajectories exhibited periods of transcription with three distinct global (i.e., including pauses) transcription velocities: one *fast*, one *slow*, and one *super-slow* (Fig. 4b), each lasting for at least tens of seconds (Fig. S2e, f, and S3a). The slow and super-slow regions were about 5 and 50 times slower than the fast region, respectively. Additionally, the super-slow regions exhibited less than half the processivity (both in terms of the number of base pairs transcribed before tether break/detachment and in terms of the number of base pairs transcribed before recovery from inhibition) compared to the slow region (Fig. S3b). Interestingly, changing the direction of the applied force did not significantly alter the interconversion rate between these regions (Fig. S3b, c).

**Figure 4:**
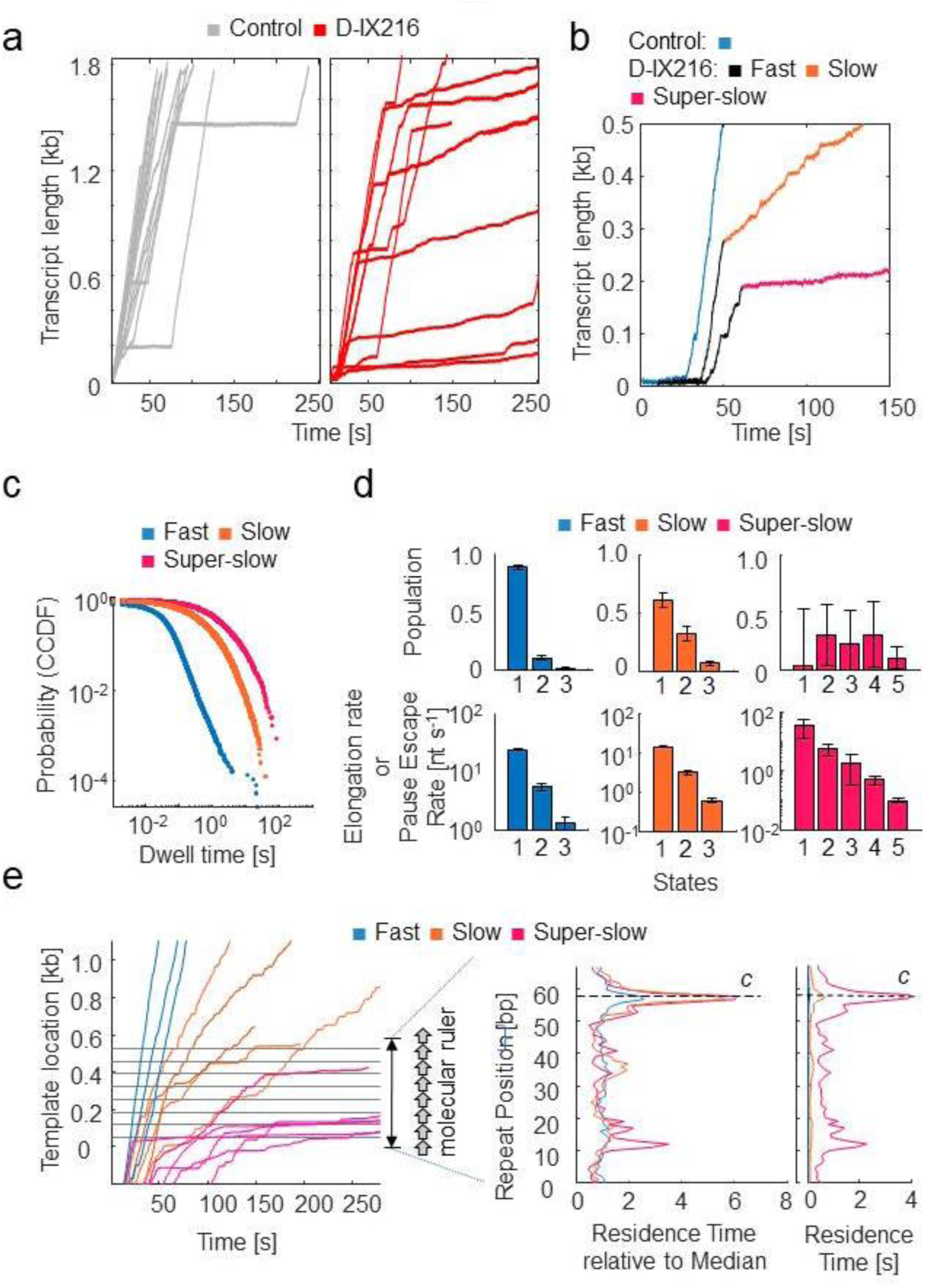
The inhibition of MtbRNAP by D-IX216 involves the conversion to one of two slowly elongating inhibited. states a) Example traces of MtbRNAP transcribing in the absence (grey) and presence (red) of D-IX216 are shown. b) Traces exhibiting the three regions, fast, slow, and super-slow, are shown and the regions are colored black, orange, and red, respectively. Transcription in the absence of D-IX216 is shown in blue. The traces plotted in this figure were collected at 18 pN of assisting force, 1 mM rNTPs, and 140 nM D-IX216. c) Comparison of the complementary cumulative distribution function (CCDF) of the dwell time distributions (DTD) of the fast, slow and super-slow regions. d) DTD analysis of the three regions is shown. e) (left) MtbRNAP traces crossing the Mtb molecular ruler are shown in varying states of inhibition. (right) The RTHes of these inhibited states are shown both in seconds or normalized to the median residence time.

The fast region resembled the enzyme’s behavior in the absence of D-IX216. DTD analysis of this region yields values similar to D-IX216-free MtbRNAP, further suggesting that the fast region corresponded to a D-IX216-free enzyme (Fig. S4). In contrast, the slow and super-slow regions were exclusively observed with D-IX216 present, indicating that they arise from its binding to the RNAP. We shall refer to the slow and super-slow regions as *inhibited states* of the enzyme, to differentiate them from the nucleotide addition states obtained from DTD analysis mentioned earlier. When bound to D-IX216, MtbRNAP displayed a consistent slow nucleotide addition rate lasting tens of seconds (Fig. 4a, b). The inhibited states persist for several steps and can interconvert with each other (Fig. S3b, c). DTD analysis of these inhibited states revealed that the slow and super-slow inhibited states consist of multiple (nucleotide addition) states, each exhibiting distinct nucleotide addition rates (Fig 4 c, d). We emphasize here that inhibited states refer to a persistent activity of the enzyme lasting for multiple incorporation cycles, while inhibited states manifest as distinct nucleotide addition dwell times. The rates (*k*’s) associated with these addition states in the fast and slow addition states differ (Fig 4d). Since there is no evidence to suggest that these addition states in the presence of D-IX216, are related to the addition states observed without D-IX216, and since their rates do not coincide, we conclude that these addition states are distinct from one another.

When the concentration of D-IX216 was increased from 70 nM to 280 nM, the kinetic parameters extracted from the DTD analysis did not change (Fig. S5a), suggesting that the slow and super-slow inhibited states occur during the prolonged binding of a single D-IX216 molecule. This inference was supported by our washing experiments, which revealed that removing D-IX216 from the transcription medium of the affected polymerase did not immediately restore normal enzyme activity (Fig. S2f). If this interpretation is correct, the conversion to the slow region results from a binding event, while reversion to the fast region is due to unbinding. Using the lifetimes of the fast and slow regions, we calculated a *k_on_*, *k_off_*, and *K*_D_ for D-XI216, obtaining 0.13 ± 0.01 µM^-1^s^-1^, 0.0032 ± 0.0014 s^-1^, and 24.6 ± 10.8 nM, respectively (Fig. S5b). The *K*_D_ so attained is consistent with the IC_50_ of ∼100 nM. We have no explanation for what process is involved in the interconversion between the slow and the super-slow inhibited states, but we speculate that it may correspond to two distinct binding modes of the inhibitor to the enzyme, affecting its activity differently.

We also examined whether the induced slowing by D-IX216 could be due to reduced rNTP affinity by decreasing the rNTP concentration from 250 µM to 25 µM (∼5x K_M_ to ∼0.5x K_M_ for the uninhibited polymerase) in the presence of D-IX216. However, the DTD behavior of the slow and super-slow inhibited states remained unchanged (Fig. S5c), suggesting that rNTP binding is not rate-limiting in these inhibited states, unlike what is expected at these concentrations in the absence of the inhibitor.

Lastly, we investigated the effects of D-IX216 on transcription across the Mtb molecular ruler to determine if these effects were sequence-dependent. Our findings revealed that slow or super-slow events occurred randomly along the DNA template, consistent with our observations using the original sequence of Mtb DNA (Fig. 4a). When MtbRNAP is in the slow inhibited state while crossing the repeats region, we observed that D-IX216 increases the residence time at every position (Fig. 4e). We normalized the RTH values to the median residence time of each inhibited state to account for the significantly different incorporation times associated with the fast, slow, and super-slow inhibited states. (Fig. 4e). This normalization demonstrated that the relative strength of the designated pause increased, as indicated by the heightened RTH peak (Fig. 4e). If pause *c* is an off-pathway pause, a kinetic competition should exist between forward translocation and pausing. Therefore, we attribute the increased strength of this pause to a decrease in the transcription rate caused by the inhibitor. In conclusion, we found that D-IX216 globally slows MtbRNAP translocation in a sequence-independent manner.

### Streptolydigin induces backtracking in MtbRNAP

Using the Mtb molecular ruler, we added 4.5-45 µM Stl to a transcribing MtbRNAP under 18pN of assisting force. We observed that Stl induced long pauses both inside and outside the repeat region (Fig. 5a). The probability of entering these long pauses (lasting over 1 second) increased from 0.42% without the inhibitor to 0.76% with Stl. Of the enzymes that recovered, the long pause consists of two components; first, the RNAP pauses with an average duration of 10 ±1.3 s (95% CI); then, it backtracks by 7 ± 1 nt (s.e.m.) and spends an additional 32 ± 4 s (95% CI) paused before transcription resumes (Fig. 5b). In contrast, previous studies have shown that Stl does not affect the backtracking dynamics of EcoRNAP^30–32^.

**Figure 5:**
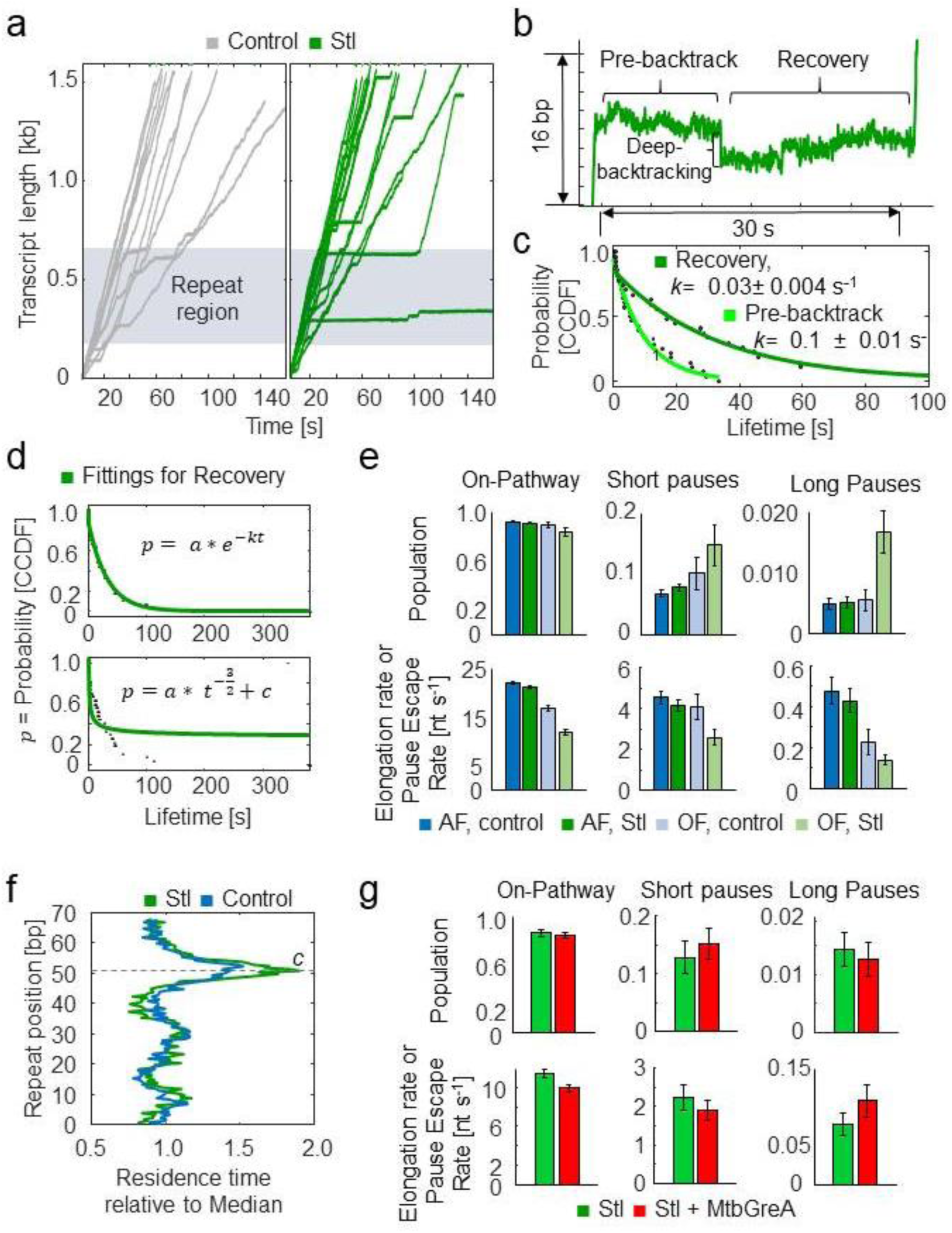
Streptolydigin inhibits MtbRNAP by enhancing pausing and inducing backtracking during pausing. a) Example traces of MtbRNAP transcription in the presence of Stl are shown without (gray) and with (green) Stl in the presence of saturating rNTP concentrations. b) Details of a backtracking event induced by Stl are shown, exhibiting a process where there is both an initial pre-backtrack pause, a backtrack, and then recovery from that backtrack. c) Fittings for the CCDFs of the pre-backtrack and recovery times of Stl-induced pauses to single exponentials are shown. d) The comparison between the fitting of the backtrack recovery between two models is shown: a single-exponential distribution corresponding to deep backtracking and a power law distribution corresponding to shallow backtracking. e) The kinetic parameters obtained from DTD analysis in the presence of Stl are shown, under a constant 18 pN assisting or 4 pN opposing forces. f) Average RTHes of MtbRNAP transcribing the repeat of the molecular ruler in the absence and presence of Stl is shown. g) Kinetics parameters extracted from DTD analysis of MtbRNAP traces under constant 5 pN of opposing force and 15 µM Stl in the absence of presence of 0.5 µM MtbGreA are shown.

We also conducted a real-time bulk fluorescence assay, during which a fluorescent nucleotide is engineered into the template strand, allowing us to distinguish between the post-translocated and the other states of the enzyme (pre-translocated or otherwise). Before the MtbRNAP arrives at this location (the pre-translocated state), the fluorescence is quenched by the base-pairing of the fluorescent nucleotide to the non-template strand. When the polymerase transits from the pre-translocated state to the post-translocated state, the fluorescent nucleotide is flipped into the active site and its fluorescence increases because its cognate rNTP is not present in solution (see Fig S6a). We used this assay to monitor the translocation state of RNAPs without cognate nucleotide and in the presence of 7.5 µM Stl^51,52^. These experiments were conducted without pyrophosphate to prevent the inherent editing of the enzyme in the backtracked state. In the absence of Stl, the MtbRNAP is predominantly found in the post-translocated state, while in the presence of Stl, only ∼20% of RNAPs remained in the post-translocated state (Fig. S6b). This reduction in the amount of post-translocated MtbRNAPs with Stl is consistent with the increase in backtracking observed in the single molecule traces (Fig 5b).

The recovery from the Stl-induced pauses before and after backtracking both exhibited single-exponential distributions (Fig 5c). A single-exponential distribution of backtrack durations is associated with *deep backtracking* and recovery via endonucleolysis,^43^ whereas a power law distribution (*t^-^*^3^*^/^*^2^) corresponds to diffusive recovery from shallow backtracking^14,16^ (Fig. 5d). Consequently, the single-exponential recovery distribution observed here primarily reflects deep backtracking, while the power law distribution also effectively describes shorter events (Fig. 5d).

Next, we compared the effect of Stl on the DTD under assisting and opposing forces. We found that Stl did not affect the MtbRNAP pause-free velocity under assisting force (Fig. 5e). However, under opposing force, which favors the pre-translocated state of the polymerase, Stl slightly reduced the polymerase velocity and significantly increased the occurrence and duration of long pauses compared to the enzyme without the inhibitor (Fig. 5e), consistent with reports regarding Stl’s effect on EcoRNAP^30^.

Our examination of the RTHs across the molecular ruler by MtbRNAP under assisting force with and without Stl, revealed that this inhibitor increased the duration of the designed pause in the tandem repeat region by 30%, but did not introduce additional sequence-dependent pause sites (Fig. 5f). This observation suggests that Stl binding is more favorable when the enzyme is paused^29^. This inference is supported by the fact that we did not observe an increased residence time background in the presence of Stl (outside of repeat pauses).

A recent article noted that factors GreA and GreB enhanced Stl’s pausing effects on EcoRNAP under opposing force^30^. However, when we assayed MtbRNAP elongation under comparable conditions using GreA (as GreB has not been found in Mtb), we observed an insignificant decrease in long pause duration (Fig. 5g). These seemingly conflicting results may reflect differences in the mechanism of Stl inhibition between EcoRNAP and MtbRNAP.

### Pseudouridimycin converts MtbRNAP into two slowly elongating inhibited states and increases the density of pauses in EcoRNAP

When PUM was added to a transcribing EcoRNAP, pauses lasting 5.9 ± 1.8 s on average were observed (pause escape rate of 0.17 ± 0.04 s^-1^), representing 2% of the enzyme’s steps (Fig. 6a). Occasionally, we noted longer pauses lasting several hundreds of seconds. Additionally, we rarely (in less than 6% of traces) observed sections where EcoRNAPs converted to one of two regions of slower nucleotide addition (slow and super-slow). These regions resemble those caused by D-IX216 on MtbRNAP, so we shall also refer to them as inhibited states; however the enzyme was not observed to recover (Fig. 6b, c).

**Figure 6:**
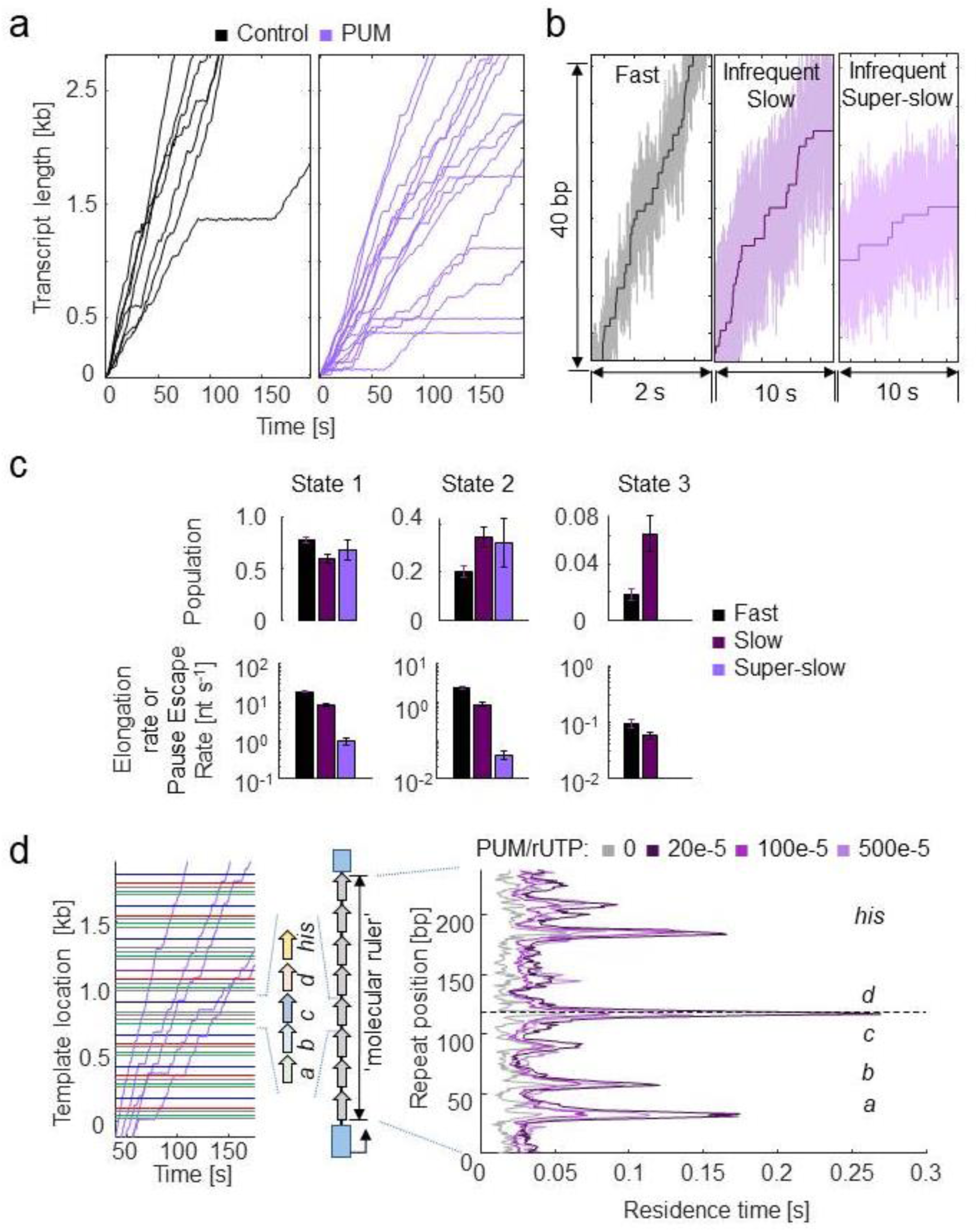
PUM inhibits EcoRNAP by enhancing pausing and infrequently inducing slowly elongating inhibited states. a) Single-molecule traces of EcoRNAP transcription in the absence (black) and presence (purple) of PUM are shown. Traces were collected at 15 pN constant assisting force, 1 µM PUM, and 1 mM rNTPs. b) A comparison of the fast region and the infrequent slow and super-slow inhibited states with PUM are shown. c) Kinetics parameters attained from DTD analysis of the three regions of EcoRNAP with PUM are shown. d) (left) Example traces of EcoRNAP crossing the molecular ruler in the presence of PUM are shown. (right) The average RTH of EcoRNAP crossing one repeat is shown at varying ratios of PUM to rUTP.

Using the molecular ruler method with EcoRNAP, our experiments showed that PUM increased the RTH peaks of the *E. coli* native pauses *his* and *d*, reduced the RTH peak of native pause *c*, and sporadically introduced more sequence-dependent pauses along the template (Fig. 6d). As PUM is an rUTP analog, it is reasonable to conclude that these additional sequence-dependent pauses occur at U incorporation sites, as it seems to be the case for the additional pausing around the *his* and *d* sites (Fig. 6d). However, due to the ± 3 bp resolution of our technique, we cannot confidently assert that all of these new peaks correspond to U incorporation sites^15^.

Interestingly, when PUM was added to MtbRNAP, a higher percentage of RNAPs (16%) exhibited conversions to slow and super-slow inhibited states reminiscent of D-IX216 binding (Fig. 7a). However, unlike with AAP, we did not observe any interconversions between the slow and super-slow inhibited states with PUM. The DTD analysis indicated that the slow inhibited states induced by PUM were composed of nucleotide addition states that were slower than wild-type (Fig 7b). Comparing the addition states from the slow inhibited states induced by D-IX216 and PUM, we found differences in both global velocity and addition states, similarly so for the super-slow inhibited states (Fig. 4d and Fig. S7a, b). Furthermore, in the presence of PUM, the polymerase displayed erratic behaviors, including the enzyme “hopping” back and forth in 10–20 bp increments, moving steadily backward (which differs from traditional backtracking), and pausing for extended periods, sometimes for hundreds of seconds (Fig. S7c, d). PUM had minimal effect on the RTH of the repeats, exhibiting only a slight increase in the strength of pause *c*, which we attribute to the inhibitor’s preference for binding to already paused polymerases (Fig. 7c).

**Figure 7:**
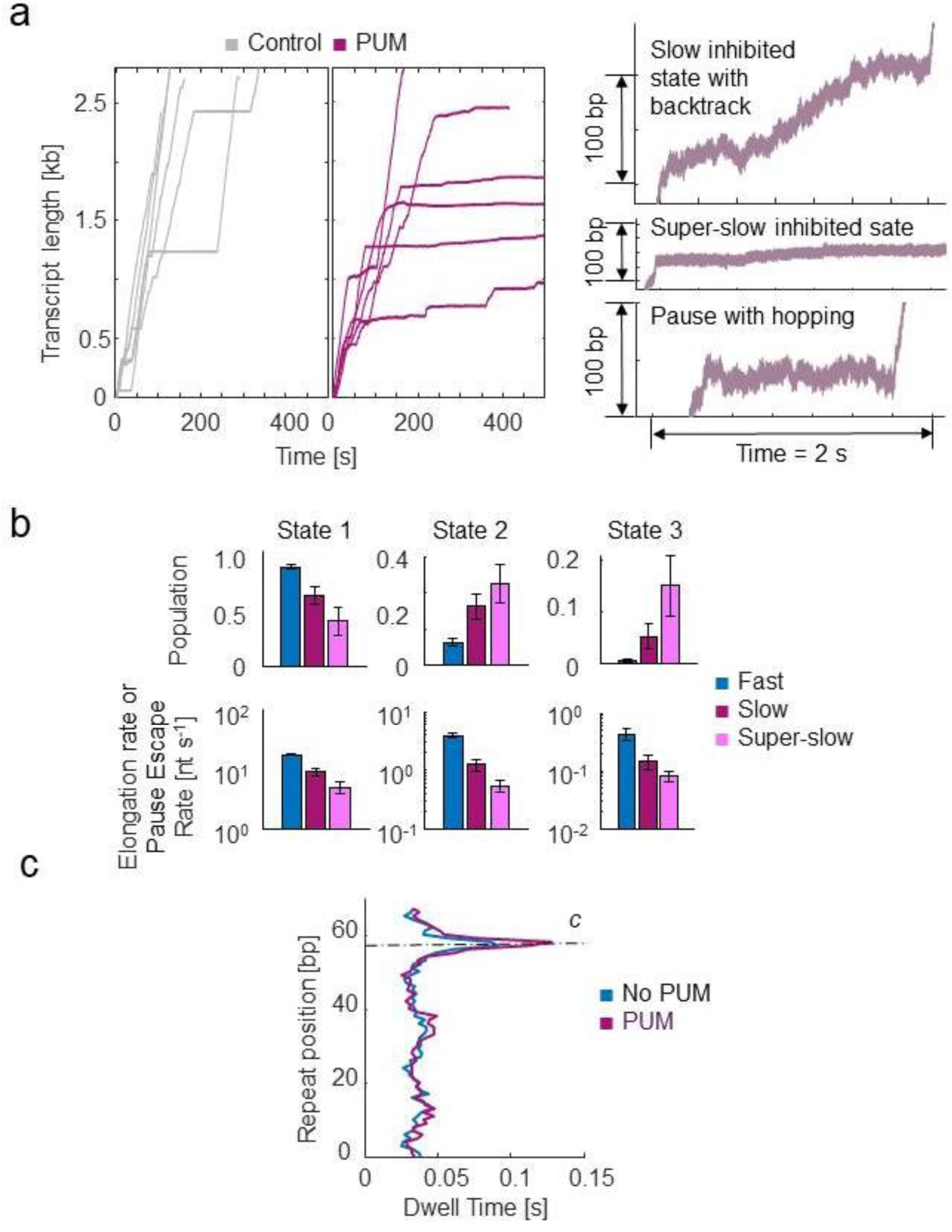
Inhibition of MtbRNAP by PUM involves induction of slowly elongating inhibited states. a) (left) Single-molecule MtbRNAP transcription in the absence (grey) and presence (purple) of PUM are shown. (right) Examples of transcription in the slow inhibited state, the super-slow inhibited state, and a dynamic pause are shown. b) Kinetic parameters attained from DTD analysis of the three regions of MtbRNAP with PUM are shown. c) The average RTH of a repeat of MtbRNAP transcribing the Mtb molecular ruler is shown.

### Mixing of Stl and D-IX216 shows a synergistic effect on transcription arrest

Treatment of tuberculosis usually involves a combination of antibiotics^53^. Accordingly, we monitored transcription by MtbRNAP in the presence of both Stl and D-IX216. Notably, Stl and D-IX216 both bind to the bridge helix of the polymerase, in distinct pockets^26,29,32,36^, which suggests that they may affect each other’s binding.

When 560 nM of D-IX216 and 15 µM of Stl were added to a transcribing MtbRNAP, the enzyme displayed either long pauses or a terminal arrest following a period of slow transcription (Fig 8a, b). The average duration of these long pauses, defined as those lasting over 5 s, was 25.2 ± 5.1 s (s.e.m.) in the presence of both inhibitors. This duration is consistent with the distribution of pause durations of MtbRNAP with 15 µM Stl alone (27.5 ± 6.7 s, s.e.m.). This suggests that these long pauses may be attributed to Stl.

**Figure 8:**
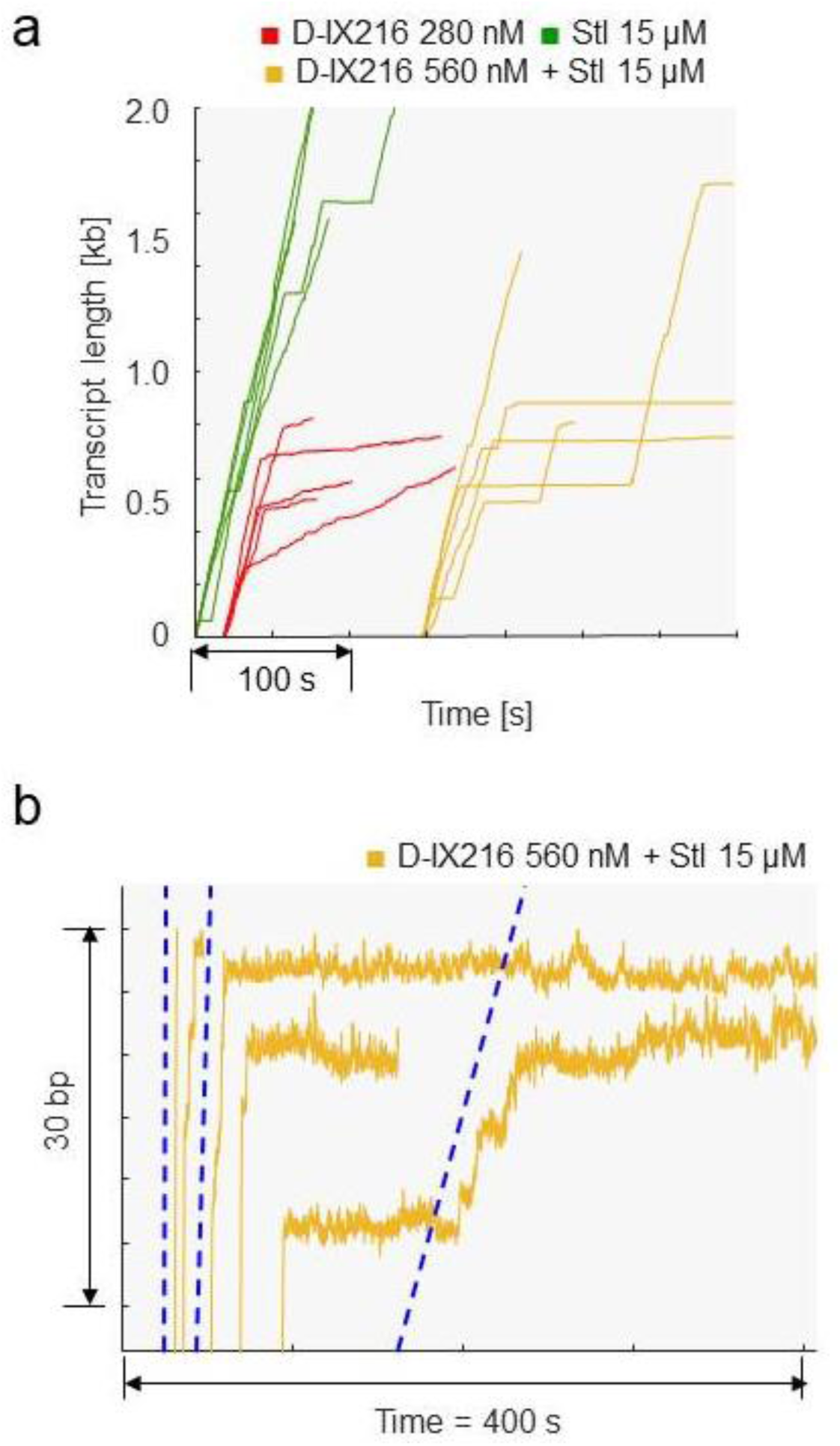
Inhibition of MtbRNAP by a combination of D-IX216 and Stl. a) Comparison of transcription elongation traces of single-molecules of MtbRNAP obtained with D-IX216, Stl, or both are shown. b) A zoom of the traces with D-IX216 and Stl just before arrest are shown. The blue dotted guide lines match the average velocity of MtbRNAP in the fast, slow, and super-slow inhibited states.

Interestingly, the arrest events following periods of slow transcription were not observed with either Stl or D-IX216 alone. The velocities during these slow sections appear to correspond to the slow and superslow inhibited states induced by D-IX216 alone (Fig 8b). Therefore, we propose that these slow transcription periods reflect the binding of D-IX216, which transitions the RNAP into the slow or super-slow inhibited state). Subsequently, the binding of Stl leads to a permanent pause (arrest).

Interestingly, while on average it takes 60 s for an Stl molecule to bind to an actively transcribing MtbRNAP at the concentration of the inhibitor used, in the presence of D-IX216, pauses arise within ∼10 s after transcription slows down. This suggests that Stl binds more readily to D-IX216-bound MtbRNAP (indicating an increased *k_on_*) than to the transcribing enzyme alone. This observation aligns with previous findings that Stl prefers to bind to already paused polymerases^29^.

In conclusion, our evidence indicates that the binding of D-IX216 can increase the *k_on_* of Stl, leading to the arrest of the enzyme. This suggests a synergistic effect between these two inhibitors, converting the lethargic nucleotide addition activity observed with D-IX216 alone into complete transcription arrests.

## Discussion and Conclusion

Our findings indicate that the global transcription velocity of MtbRNAP is slower than that of EcoRNAP, which mirrors the trend in total transcription rates observed *in vivo* ^54^ In addition, whereas MtbRNAP has comparable mechanical robustness to EcoRNAP, it generally possesses weaker pausing. Our molecular ruler and bulk data suggest that the consensus pause sequences of the *E. coli* enzyme are also recognized by MtbRNAP, but significantly more weakly.

Our single-molecule studies demonstrate that three small-molecule inhibitors of MtbRNAP: D-IX216, Stl, and PUM, do not act as chain terminators, but instead slow transcription elongation in distinct ways.

D-IX216 inhibits transcription by inducing two distinct inhibited states that incorporate nucleotides more slowly. Our findings show that this inhibitor does not completely stop nucleotide incorporation, suggesting that it is engaged during elongation and exerts an inhibitory effect that differs from other inhibitors that target the bridge helix of the RNAP^26,32,36,37,51,55–58^. The periods of recovery from transcription elongation inhibition observed in our molecular trajectories likely correspond to unbinding events of the inhibitor. The different nucleotide addition states, associated with different rates of nucleotide incorporation, may reflect stochastic differences in the functional interactions between the catalytic elements of the RNAP and the inhibitor. Future research should investigate the causes of the inhibited states.

Previous studies have shown that Stl lengthens existing pauses, which is consistent with our observation that pause *c* is extended with Stl. However, those studies did not explicitly describe the effect of Stl on backtracking^30^. We find that Stl induces pausing in RNAP, followed by backtracking. This finding may be attributed to the superior capabilities of single-molecule methods over bulk methods, particularly in terms of spatiotemporal resolution.

PUM increases the pause density in both EcoRNAP and MtbRNAP by competing with UTP incorporation^35^. In addition, PUM can transform the active polymerase into two distinct slowly-elongating inhibited states that do not appear to interconvert. It is possible that the inhibited states induced by AAP and PUM share similar underlying mechanisms of inhibition, but this cannot be determined definitively, given that the elongation rates of the two inhibited states differ for the two inhibitors.

Both Stl and D-IX216 bind to the bridge helix of MtbRNAP. When administered together, D-IX216 binding seems to promote the binding of Stl, resulting in more potent inhibition (or arrest) of the enzyme. Since the standard of care for the treatment of tuberculosis infection involves a combination of antibiotics, identifying inhibitors that work together effectively is particularly valuable, and is not routinely the case^59^. The results presented here with the combination of Stl and D-IX216 provide a basis for their possible use in combination therapy. Developing distinct antibacterial compounds that act differently or have diverse mechanisms of inhibition of MtbRNAP is essential to overcome the challenges of drug resistance, develop reduced duration of treatments, and improve patient outcomes.

## Materials, Data, and Code Availability Statement

All oligos/plasmids used in this study are available upon reasonable request. See Supplementary Tables S1-S3 for a list of materials. Transcription traces are available from Dryad at https://doi.org/10.5061/dryad.2fqz6130m (**Private peer review preview at http://datadryad.org/stash/share/I7CMyC4-ak9FzzgBQz0WtYGq7fTjrPCZa7J6iYap2iE). Matlab code used for analysis is available on Github at https://doi.org/10.5281/zenodo.6534021.

## Acknowledgments

We would like to extend thanks to Keren Espinoza and Cristhian Cañari for their work in the early stages of this project. This research was funded by National Institutes of Health grants GM041376 to R.H.E. and R01GM032543 to C.B.. W.L. was supported by the National Natural Science Foundation of China grant 31900883, Department of Biomedical Engineering, Shenzhen University. C.B. is a Howard Hughes Medical Institute investigator.

## Methods

A list of oligos, plasmids, and cells used are available in Supplementary Tables S1-4.

### Plasmid preparation

To make the vectors for MtbRNAP biotinylation, we performed round-the-horn site-directed mutagenesis of plasmid pET-DUET-BC^60^ using long PCR to introduce an avitag or a sortase tag (sortag) sequence at the c-terminus of the rpoC gene^13^. The resulting plasmids were named pET-DUET-BC-avitag or pET-DUET-BC-sortag, respectively. As for the vectors for expressing Mtb transcription factors, we amplified the corresponding gene from the Mtb genome and inserted it into the plasmid pET His6 TEV LIC (Addgene, # 29653) using ligation-independent cloning^61^.

To make plasmids with the DNA templates for the single-molecule assays, we took pUC19 and cloned the promoter AC50, the 10-bp stalling cassette, and the transcribable region, which derived from a ∼ 2.7 kb region of genes rpoB and C from the *M. tuberculosis* Erdman genome (a gift from Professor Sara Stanley). We used an additional round-the-horn site mutagenesis to remove the lac promoter. We named this plasmid pUC19-del-Lac-AC50-MtbrpoBC. We used this plasmid to create another plasmid containing a molecular ruler. First, we removed one side of the BsaI in the rpoC gene by inserting a 255 bp DNA fragment of terminator WhiB1. Second, we replaced the stalling sequence with a 20-bp CU-less cassette using round-the-horn mutagenesis. Third, we used NEBuilder® HiFi DNA Assembly Master Mix (NEB E2621S) to insert a synthetic DNA fragment, an 8x repeat of a 63 bp sequence of pause *c* (Genescript), into the rpoB gene. The resulting plasmid was named pSGUML1T1.

### DNA template preparation

We made a single-end digoxygenin-labeled DNA template (without a ruler) using PCR by amplifying approximately 4.2 kb or 2.8 kb from plasmid pUC19-del-Lac-AC50-MtbrpoBC, depending on whether the assay is for opposing or assisting force, and utilized a dig-labeled primer.

To make the DNA template containing the molecular ruler, we linearized the plasmid pSGUML1T1 with BsaI and filled the sticky ends by using Klenow Fragment in the presence of either ddATP or ddCTP to produce the opposing or assisting force template, respectively^13^.

In addition, DNA handles were created using PCR with a biotinylated primer along with another primer, either double-dig labeled or encoding a restriction enzyme site for generating sticky ends for later ligation.

### Production of biotinylated and non-biotinylated MtbRNAP

We produced the holoenzyme of MtbRNAP with sigma factor A by co-expressing its subunits in *E. coli*^60^. We biotinylated the avitag-MtbRNAP by co-expressing the plasmids pACYD-DUET-AOS and pDUET-BC-avitag with the plasmid pCDF-BirA (a gift from Professor Nicola Burgess-Brown). This protein was used for the assay under opposing force with Stl and GreA. Alternatively, we biotinylated the sortag-MtbRNAP (LPETG at the c-terminal of β’) after purification using an *in vitro* reaction of the polymerase, the enzyme sortase A^62^ and GGGL-biotin (Genscript)^63^.

### Production of transcription factors MtbCarD and MtbGreA

We purified the transcription factor CarD as before^60^. To obtain MtbGreA, we expressed it as a his-tagged protein in *E. coli* BL21(DE). We captured this protein with Histrap ® (Cytiva) and used a TEV protease cleavage reaction^64^ to cleave its histidine tag. To remove the cleaved tag, we used reverse nickel chromatography. The final purification step was done with anion exchange chromatography (HiTrap Q, cytiva) in a Tris-saline buffer.

### *In vitro* transcription bulk assay

We used an artificial bubble-based transcription to assess the recognition of the elemental pause sequences by MtbRNAP using a FAM-labeled RNA^60^ and assessed transcript size via gel. Raw gels are available in the Source Data Files related to the relevant figures.

### Ternary elongation complex formation

We stalled the biotinylated MtbRNAP on a cassette lacking one or two nucleotides to form the ternary elongation complex (TEC) to study elongation, as previously described^65,66^.

### Sample preparation for single molecule optical tweezers transcription elongation assay

To create the promoter-initiated stalled MtbRNAP elongation complexes, we utilized MtbCarD. First, we pre-incubated approximately 1 µM of biotinylated-MtbRNAP with around 10 µM of MtbCarD at 37 °C for 5 minutes in a final volume of 2 µL. Subsequently, we diluted the biotinylated MtbRNAP in buffer TB40 (20 mM Tris-HCl pH 7.9, 40 mM KCl, 5 mM MgCl_2_, and 0.5 mM TCEP) to a concentration of 10 nM. We then stalled 5 nM of polymerases by incubating them with 150 µM GpG, 2 nM Mtb DNA template in stall Buffer (20 mM Tris-HCl pH 7.9, 3 µM rATP, 3 µM rGTP, and 3 µM rCTP) at 37 °C for 20 minutes. If we were to use the Eco DNA repeat, we would substitute GpG with ApU^67^.

To attach the DNA handles or stall complexes to the microbead surfaces, we used a few different methods. In the first method, we incubated a 0.1 nM stalled complex labeled with double-digoxygenin with 0.05-0.1% passivated anti-dig beads in TB40 buffer for 10 minutes at 22 °C. Then, we added 0.05 µg/µL heparin at 22 °C for 5 minutes to remove non-competent polymerases. In the second method, we incubated 0.2-0.5 nM digoxygenin-labeled DNA handle with 300 nM neutravidin and the same amount of anti-digoxigenin beads at 22 °C for 10 minutes, followed by the addition of heparin. In the third method, we conducted ligation-based deposition by incubating a 0.1 nM stall complex with 0.1-0.05% passivated oligo beads, 0.1 mM rATP, and 4 U/µL T4 DNA ligase in TB40 buffer at 22 °C for 45 minutes to facilitate the ligation reaction. This was followed by a 5-minute treatment with heparin at 22 °C. In the fourth method, we attached 0.2-0.5 nM DNA handle by incubating it with passivated complementary oligo beads, 300 nM neutravidin, 0.1 mM rATP, and 4 U/µL of T4 DNA ligase in TB40 buffer for an identical prolonged incubation, followed by heparin treatment. The bead depositions were either kept on ice for immediate use, or preserved by flash freezing and stored at -80°C.

For every 10 µL of deposited beads, we added 1,000-1,500 µL of degassed TB130 Buffer (20 mM Tris-HCl pH 7.9, 130 mM KCl, 10 mM MgCl_2_, 1 mM TCEP and 5 mM NaN_3_) before injecting into the fluidics chamber.

In the experiments with MtbCarD and MtbGreA, the final amount of initiating CarD contributed to the amount free in solution in elongation was as low as ∼13 pM (diluted from 100-120 nM MtbCarD), and neither causes any effects on pausing or velocity.

### High-resolution optical tweezer transcription elongation assay

In Fig. 1a there is a representation of the optical tweezers setup, and in Figure 1b, 1c, and 2b there are representations of assisting force, opposing force, and passive modes experiments. We used a custom home-built timeshared dual-trap optical tweezers system^67,68^. In short, the single molecule assay to measure transcription elongation involves two beads, one coated with RNAP stalled complexes, and one coated with DNA handles with neutravidin on the free end. The two beads are caught in separate traps, and when the beads are brought in close proximity, the neutravidin on the end of the DNA handle binds to the biotin attached to the RNAP and a tether is formed. The beads are brought apart, and the DNA handle separates the RNAP from the surface of the bead to reduce photodamage. We also used an oxygen-scavenger system based on sodium azide to increase the robustness of collecting processive activity. For both EcoRNAP and MtbRNAP, the attachment to biotin is done via its β’ subunit.

We injected the microbeads into a microfluidics chamber through different channels to diffuse them to a central main channel, trapped them by optical tweezers and brought them close to each other from the tether. To resume the elongation activity, we flowed a solution containing rNTPs, transcription factors, and/or inhibitors through a shunt positioned very close to the tethered complex. We used fluidic valves to control the flow via gravity. When transcribing the Eco repeat sequence, we used a nucleotide cocktail of 1 mM rUTP, 1 mM rGTP, 0.5 mM rATP, and 0.25 mM rCTP^13^ to match previous studies.

The instrument records the trap separation and monitors the displacement of the beads from the centers of the traps, and handles force feedback for the constant force experiments^63^. Calibration from raw voltages from the instrument to distance (bead separation) and applied force is done via power spectrum calibration to the expected Brownian motion of a bead held in a trap^67^.

### Analysis of single-molecule complete trajectories of transcription elongation

All analysis, including the processing of the raw instrument data to forces and bead-to-bead distances and analysis procedures is done via custom code written in MATLAB, and is available at https://doi.org/10.5281/zenodo.6534021.

The collected traces are first selected for their length and undergo an initial sorting step where traces with unusually high instrumental noise or those that are unfit for analysis (two RNAPs tethered at the same time, too short transcription, etc.) are removed.

For calculating velocity distributions, a Savitzky-Golay differentiating filter of degree 1, width 1s is used to extract the velocity of the trace at every point. These velocities are binned and least-squares fit to the sum of two Gaussians, one with mean 0 bp/s.

To get the dwells of the polymerase, we fit the traces as a staircase with a 1 bp step size. The complementary cumulative distribution function (CCDF) of the dwell time distribution (DTD) is fit to a sum of exponentials (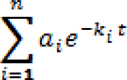), where *i* represents the *i-th* nucleotide addition state of the RNA polymerase, *a_i_* is the state probability 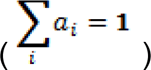, and *k_i_* is the kinetic rate such that *k_1_* > *k_2_* > … *k_n_*. The CCDF is fit to the sum of up to 6 exponentials, and the best is chosen via the Akaike information criterion.

We calculated the residence time histogram (RTH) of the polymerase transcribing the pauses repeat according to a previous publication^13^. Briefly, the repeating pauses lets us determine a scale factor and offset factor to each transcription trace based on the location of the pauses. The scale is chosen to space the pauses 68 bp apart, and the offset is chosen to place the pause at position 50 bp into the repeat. This is done by taking the RTH of the repeat, averaged across the 8 repetitions, for a range of scale factors. The one that aligns the pauses best is the one with the most peaked RTH (a poor scale factor does not align the pauses, so makes the RTH flat, a good scale factor aligns the pauses and thus it will show up as a peak). The offset factor is then chosen to place the location of the strongest peak in the best RTH at 50 bp.

A tabulation of the number of traces for each experimental condition is available in Table S5.

## Supplementary Figures

**Figure S1:**
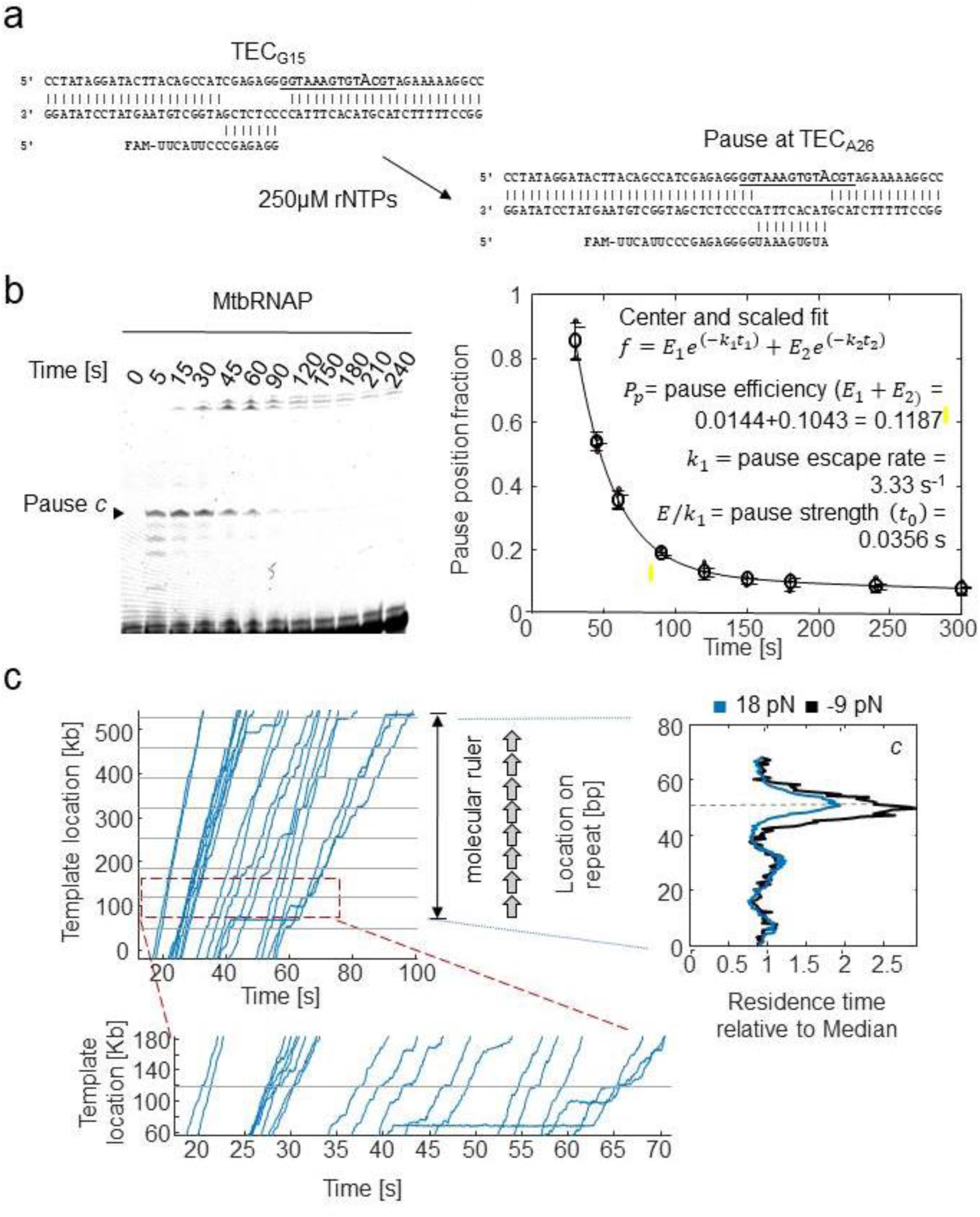
The design of a *molecular ruler* to study pausing in MtbRNAP with high-resolution optical tweezers. a) An artificial transcription bubble scheme with a 5’-end labeled RNA with FAM was used for bulk elongation assay. The stalled ternary elongation complex at position G15 (TEC_G15_) resumes RNA polymerization activity upon adding 250 µM rNTPs. b) MtbRNAP efficiently recognizes sequence derived from *E. coli* elemental pause *c*. (left) A urea PAGE gel that shows the clearance of the pause site at TEG_A26_ with a mean duration of ∼ 27 sec (inverse of the pause escape rate). (right) The quantification of the bands and their fitting to a bi-exponential decay. The extracted parameters indicate that the pause escape rate is kinetically equivalent to *his* pause of EcoRNAP^70^. c) The *Mtb* molecular ruler enables the analysis of sequence dependence pausing by MtbRNAP at the single-molecule level. Example traces are shown on the left, collected at 1mM rNTPs and 18 pN assisting force. Below is a zoom around the designed pause. The RTH on the right shows the strong pausing at 50 bp, and how the strength changes with the direction of the applied force. Figure S1—Source Data 1: Original gels for Figure S1b, indicating the conditions and relevant bands Figure S1—Source Data 2: Original files of the gels in Figure S1b.

**Figure S2:**
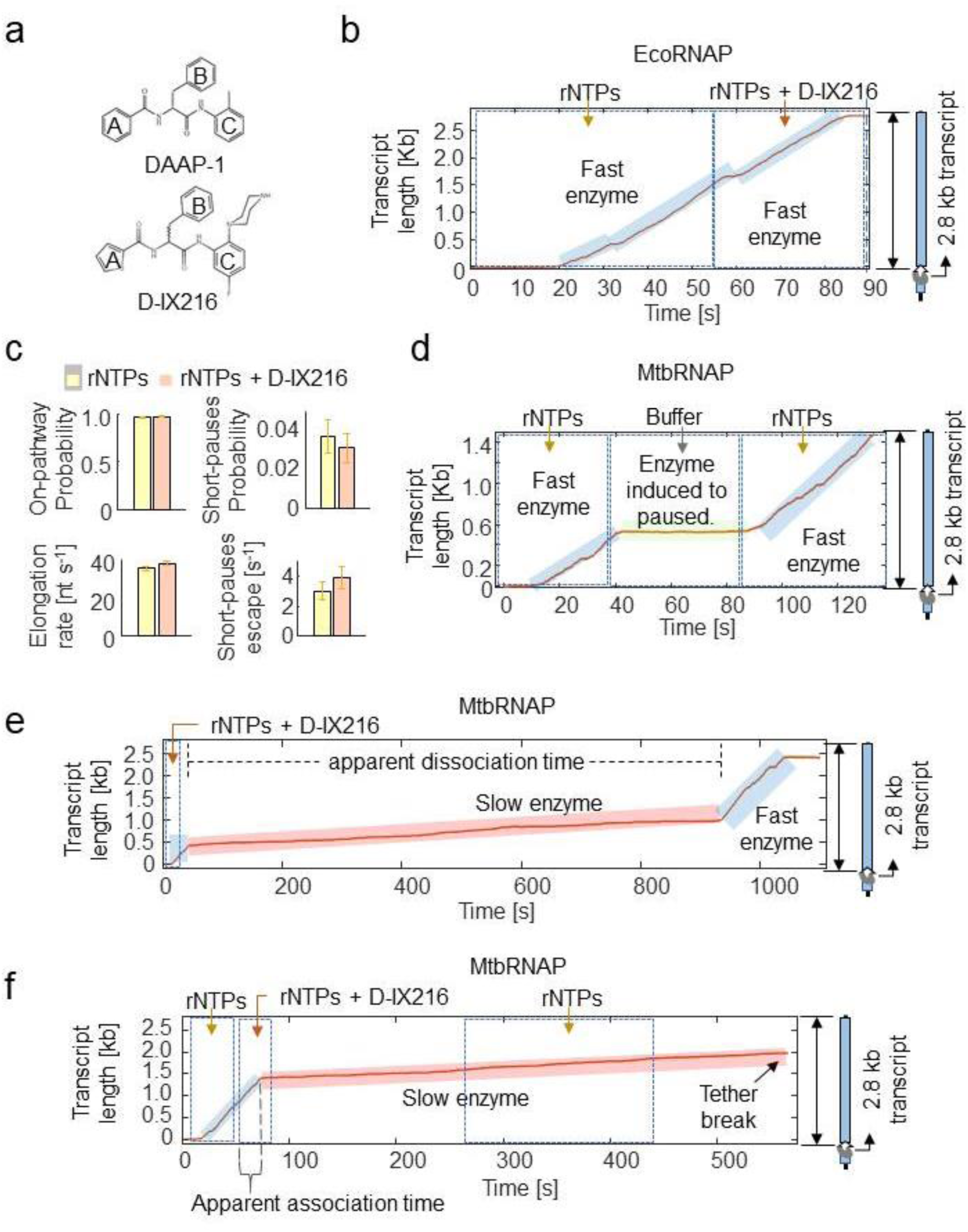
D-IX216 specifically slows down the MtbRNAP transcription. a) The chemical representation of the Nα-aroyl-N-aryl-Phenylalaninamides (AAPs) D-AAP1 and D-IX216 are shown. Rings A and C are modified to increase the affinity of D-IX216 for MtbRNAP^25^. b) D-IX216 does not alter a single EcoRNAP activity. An example trace of a two-shunt experiment shows that 280 nM D-IX216 does not affect EcoRNAP activity. The dotted boxes denote what solution is flowing out of the shunt at what time. c) The kinetic parameters obtained from DTD analysis of EcoRNAP in the absence and presence of D-IX216 is shown. In this dataset, there were too few events to fit a third exponential (long pauses). d) This buffer exchange control experiment shows that the RNAP quickly responds to buffer changes. When nucleotides are removed, MtbRNAP immediately pauses until the nucleotides are reintroduced. e) D-IX216 slows down MtbRNAP rather than pausing or halting it. The trajectory of a single MtbRNAP elongating at constant 18 pN assisting force with 280 nM D-IX216 and saturating nucleotides (∼1 mM) shows a slow state that eventually instantly recovers its velocity (∼ 20 bp/s). Depending on the condition, up to ∼ 41% of polymerases can recover their initial velocity. The dotted boxes show the start and duration of the shunt opening to supplement the inhibitor and nucleotides. The apparent association time (*t_a_*) for D-IX216 binding corresponds to the time between the shunt opening to introduce D-IX216 and the switch’s appearance. On the other hand, the apparent D-IX216 dissociation time corresponds to the slow region’s lifetime. f) D-IX216 is still engaged in MtbRNAP elongation. We performed a buffer exchange using two shunts to remove the inhibitor from the affected polymerase. However, this did not immediately restore its global velocity. Towards the end of the trace, the polymerase might detach from the DNA template, leading to tether break events or stalls in 45% and 13% of the polymerases, respectively.

**Figure S3:**
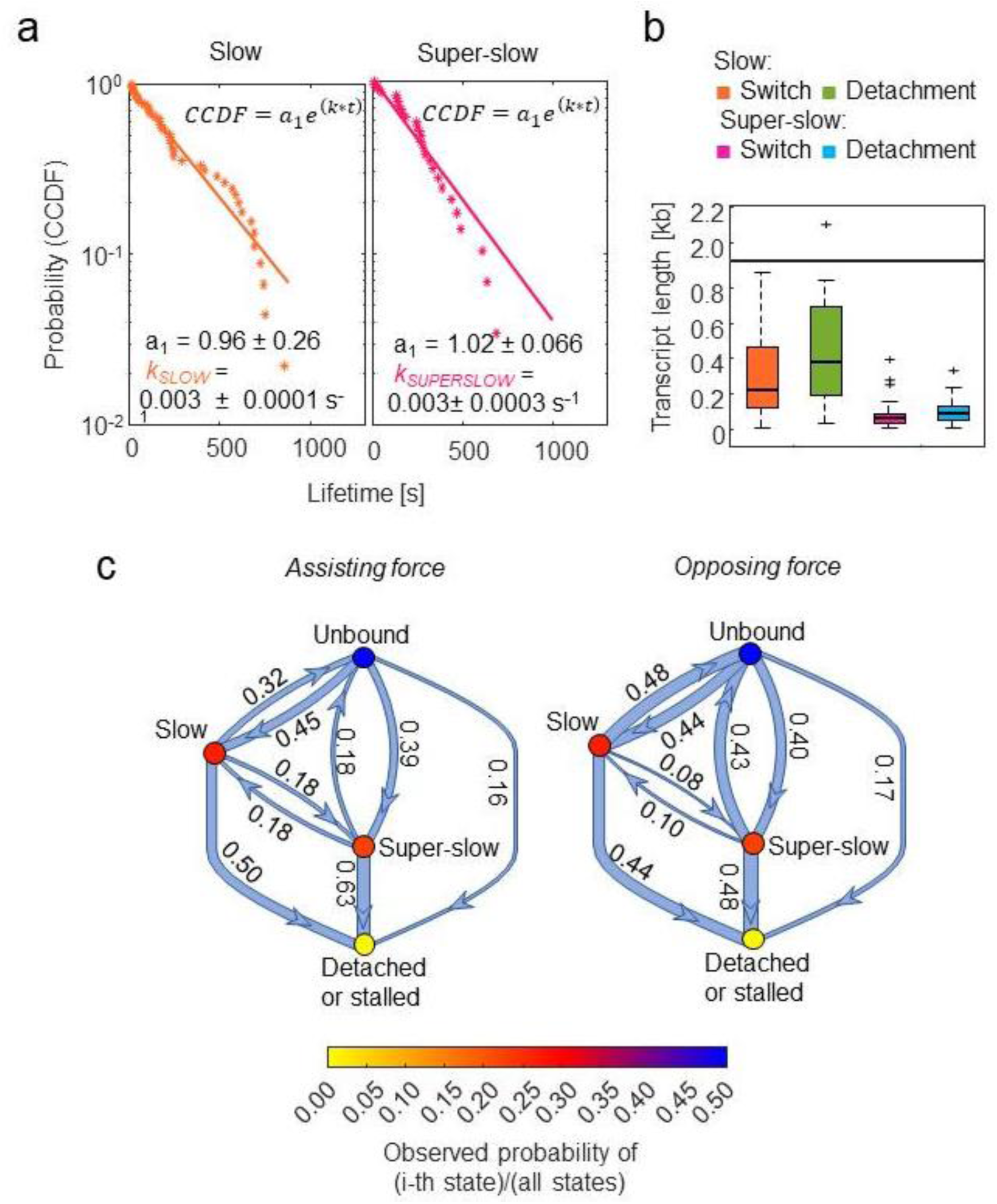
Interconversion between the slow and super-slow inhibited states. a) The CCDFs of lifetimes of the slow and super-slow inhibited states are shown with a fitting to a single exponential decay. The lifetime was measured as the time between switching into that state, until the motor either recovered (fast) or switched into the other state (e.g. a switch from slow to super-slow). Tether-break events were not included. b) Comparison of the processivity (in terms of kb extended by the elongating polymerase before tether break (template dissociation), or in terms of kb before a switch event of the super-slow and slow inhibited states is shown. Processivity is lower in the super-slow inhibited state than in the slow inhibited state by both definitions. The data for the bar plots were obtained from combined data of assisting and opposing forces experiments. c) Transition maps under assisting and opposing force of the different states (fast, slow, and super-slow) are shown. Opposing force increases the probability that the enzyme recovers to the fast (unbound) state compared to assisting force.

**Figure S4:**
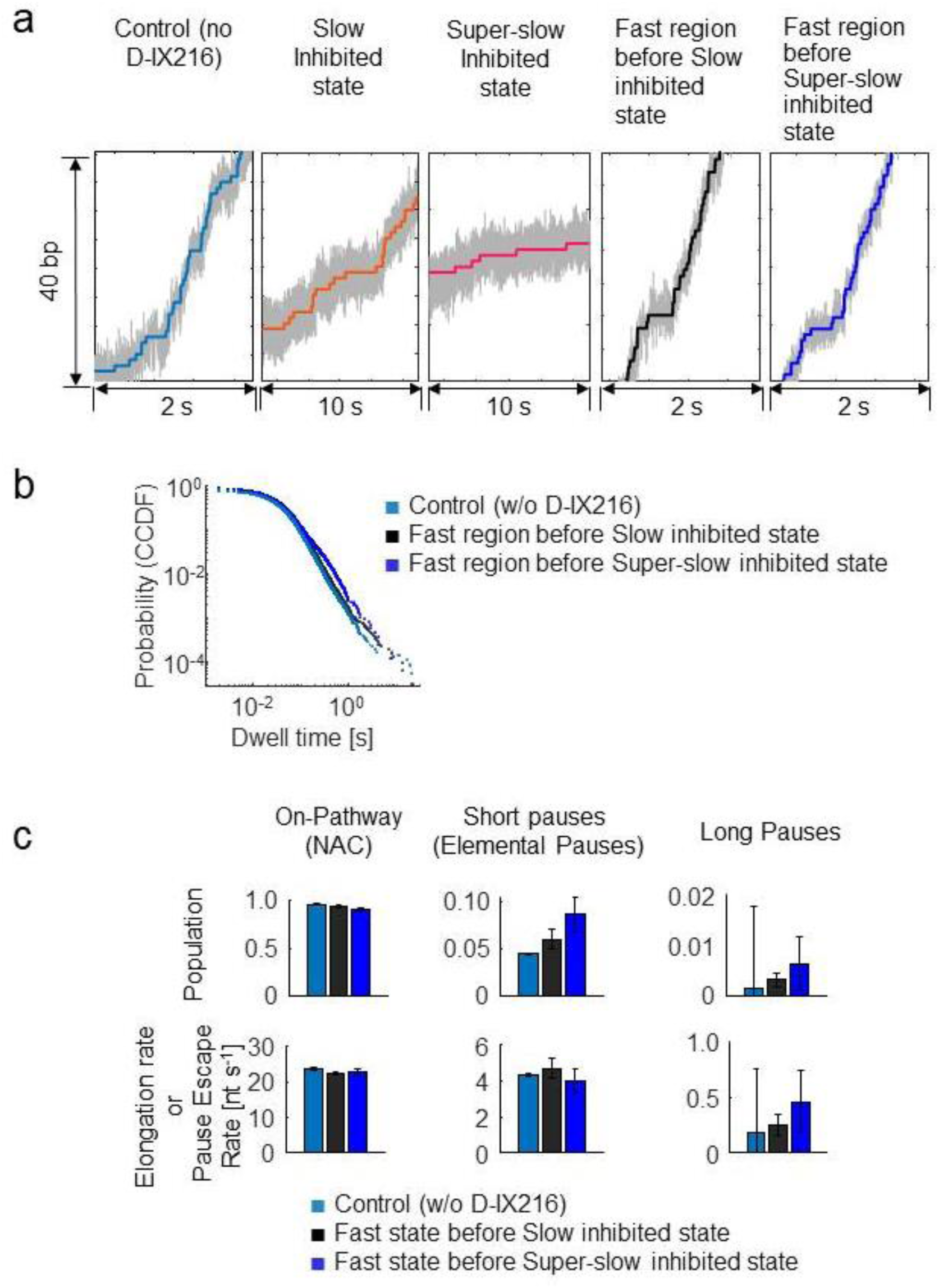
The fast elongating state of MtbRNAP observed in the presence of D-IX216 corresponds to the inhibitor-free state. a) Example fittings of the staircase stepping fitting for different inhibited states of MtbRNAP are shown. b) The CCDFs of the DTDs for fast elongating polymerases observed with (black and dark blue) and without D-IX216 (cyan). c) The kinetic parameters from DTD analysis for the enzyme without D-IX216 and the enzyme in the fast state with D-IX216 are shown. No notable differences are found between the inhibitor-free and the fast state with D-IX216.

**Figure S5:**
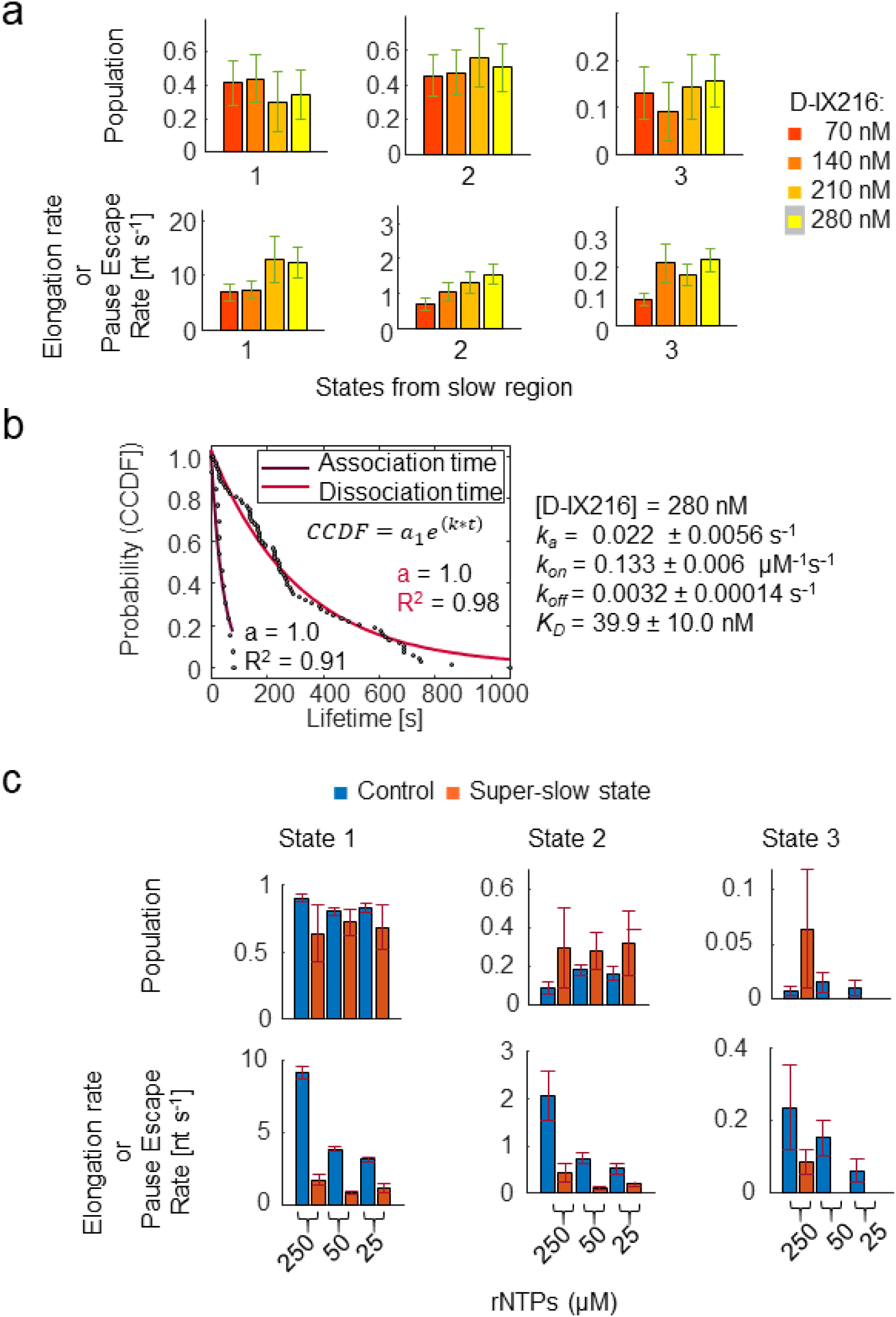
Effect of different D-IX216 and nucleotide concentrations on the slow and super-slow inhibited states of MtbRNAP. a) The concentration of D-IX216 did not impact the kinetic parameters obtained by DTD analysis of MtbRNAP in the slow inhibited states. This suggests that a single molecule of D-IX216 causes the conversion into the slow state. b) Measurement of the dissociation constant (*K_D_*) of D-IX216 for the elongating MtbRNAP. The association and dissociation times obtained from the duration of the fast, slow, and super-slow inhibited states in the single-molecule traces are fit to single-exponentials to obtain k_on_ and k_off_. Their ratio is K_D_. c) The concentration of rNTPs did not affect the kinetic parameters in the slow inhibited state of MtbRNAP induced by D-IX216. This indicates that the slowing by D-IX216 is not due to a reduction in nucleotide affinity. These experiments were done in semipassive assisting force mode, where the optical traps are moved stepwise to keep the force within a set range. In other words, the traps are held at constant position, causing the force to fall as transcription progresses, until it reaches some lower force limit (in this case, 10pN) after which the traps move apart to raise the force to some upper force limit (in this case, 18pN).

**Figure S6:**
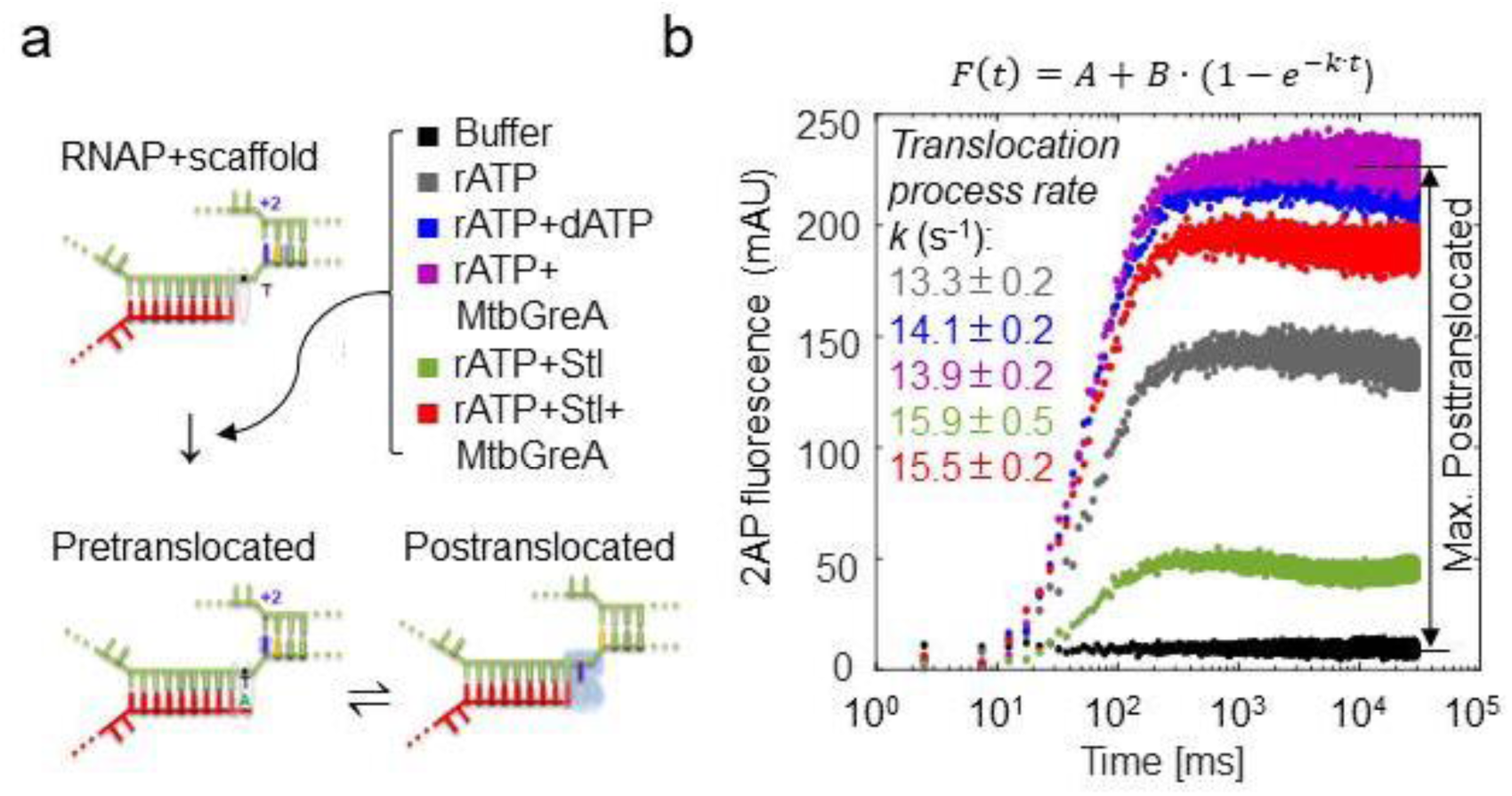
Streptolydigin (Stl) induces reduced post-translocated states in MtbRNAP. a) A bulk assay was performed to measure the translocation of RNAP by one nucleotide using the Mtb ternary elongation complex. The technique involved real-time measurement of the fluorescent enhancement at 375 nm of a nitrogenous base analog, 2-aminopurine^52^. This analog was positioned on the template strand of the DNA-RNA hybrid, initially at position +2, and then moved towards position +1 due to the RNAP translocation process. The workflow included the use of a fluorescent artificial bubble to measure the effect of Stl on the translocation state of MtbRNAP. After the RNAP incorporated one rATP, the translocation state was measured by the dye’s fluorescence in the following base in the template DNA. When the base was paired, fluorescence was low due to quenching; when the base was unpaired, such as in the post-translocated state, the fluorescence was high. The assay conditions consisted of transcription buffer and 100 µM rATP, along with either 100 µM dATP, 0.25 µM MtbGreA, 7.5 µM Stl, or 0.25 µM MtbGreA / 7.5 µM Stl. b) The comparison of different conditions of the translocation trace of the MtbRNAP in the presence of cognate nucleotide (rATP) and Stl shows that the inhibitor allows only ∼ 20% of the MtbTEC to attain the post-translocated state (green dots). The addition of MtbGreA was used as a positive control for post-translocation.

**Figure S7:**
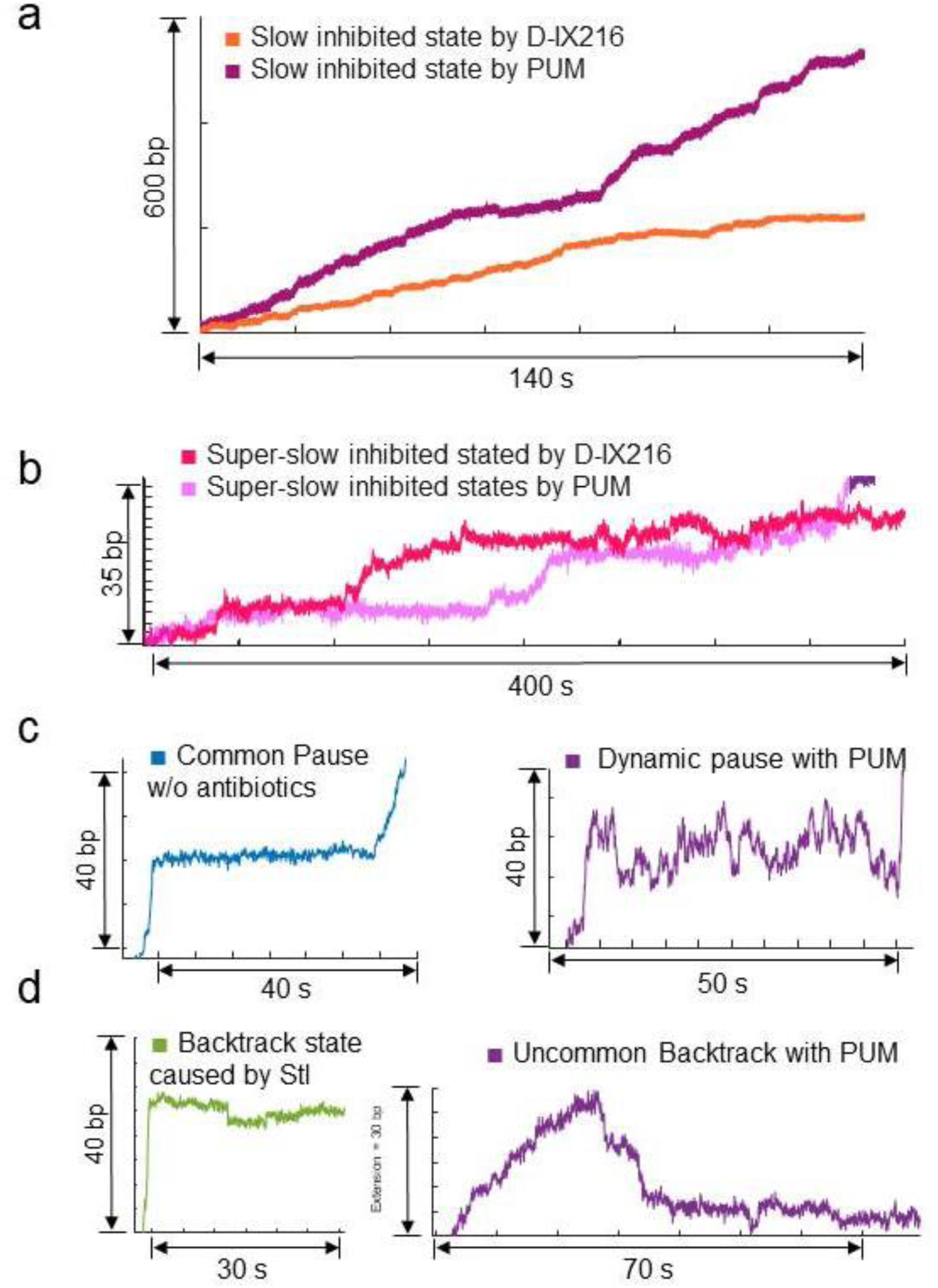
Comparison of events found in transcription with antibiotics. a) Example traces in the slow inhibited state caused by PUM and D-IX216 are shown. b) Example traces in the super-slow inhibited state caused by PUM and D-IX216 are shown. c) Comparison of a common pause obtained in the absence of inhibitor versus a dynamic pause caused by PUM is shown. d) Comparison of a backtrack event observed in the presence of Stl compared to a rare non-traditional backtrack event induced by PUM is shown.

